# The development of cooperative channels explains the maturation of hair cell’s mechanotransduction

**DOI:** 10.1101/654764

**Authors:** Francesco Gianoli, Thomas Risler, Andrei S. Kozlov

## Abstract

Hearing relies on the conversion of mechanical stimuli into electrical signals. In vertebrates, this process of mechano-electrical transduction (MET) is performed by specialized receptors of the inner ear, the hair cells. Each hair cell is crowned by a hair bundle, a cluster of microvilli that pivot in response to sound vibrations, causing the opening and closing of mechanosensitive ion channels. Mechanical forces are projected onto the channels by molecular springs called tip links. Each tip link is thought to connect to a small number of MET channels that gate cooperatively and operate as a single transduction unit. Pushing the hair bundle in the excitatory direction opens the channels, after which they rapidly reclose in a process called fast adaptation. It has been experimentally observed that the hair cell’s biophysical properties mature gradually during postnatal development: the maximal transduction current increases, sensitivity sharpens, transduction occurs at smaller hair-bundle displacements, and adaptation becomes faster. Similar observations have been reported during tip-link regeneration after acoustic damage. Moreover, when measured at intermediate developmental stages, the kinetics of fast adaptation varies in a given cell depending on the magnitude of the imposed displacement. The mechanisms underlying these seemingly disparate observations have so far remained elusive. Here, we show that these phenomena can all be explained by the progressive addition of MET channels of constant properties, which populate the hair bundle first as isolated entities, then progressively as clusters of more sensitive, cooperative MET channels. As the proposed mechanism relies on the difference in biophysical properties between isolated and clustered channels, this work highlights the importance of cooperative interactions between mechanosensitive ion channels for hearing.

**SIGNIFICANCE:** Hair cells are the sensory receptors of the inner ear that convert mechanical stimuli into electrical signals transmitted to the brain. Sensitivity to mechanical stimuli and the kinetics of mechanotransduction currents change during hair-cell development. The same trend, albeit on a shorter timescale, is also observed during hair-cell recovery from acoustic trauma. Furthermore, the current kinetics in a given hair cell depends on the stimulus magnitude, and the degree of that dependence varies with development. These phenomena have so far remained unexplained. Here, we show that they can all be reproduced using a single unifying mechanism: the progressive formation of channel pairs, in which individual channels interact through the lipid bilayer and gate cooperatively.

## INTRODUCTION

Mechano-electrical transduction (MET) is a fundamental process that transforms auditory stimuli into electrical signals that propagate through the brain. In vertebrates, this transformation takes place when a mechanical stimulus deflects the stereocilia of sensory receptors in the inner ear, the hair cells, opening mechanosensitive ion channels (1, 2). Stereocilia are actin-filled, enlarged microvilli arranged in a staircase manner in a hair bundle. Each stereocilium connects to its taller neighbor by a filamentous linkage, the tip link, necessary for mechanotransduction (3).

Tip links are elastic elements that tense or relax in response to the hair bundle’s deflections and transmit force onto the MET channels, changing their opening probability (4). When a hair bundle is deflected toward its tall edge, referred to as the positive direction, the MET channels open, but then close again with time (5). This phenomenon, named adaptation, appears as a combination of two distinct processes: the first one is ‘fast’, with a time constant of a few milliseconds or even less in mammals, and the second one is ‘slow’, with a time constant of a few tens of milliseconds. Fast adaptation is accompanied by a rapid movement of the hair bundle in the direction opposite to the stimulus, called ‘the twitch’ (6, 7). It is believed to be caused by the direct reclosure of the MET channels and is at least partially dependent on the action of Ca^2+^ ions (7–10). Slow adaptation, in contrast, compels the hair bundle to move in the direction of the stimulus and affects the channels’ open probability via tip-link tension regulated by myosin motors (11).

Mechanotransduction in hair cells matures gradually before the onset of hearing (12–14). In mice and rats, cochlear hair cells are insensitive to mechanical stimuli before birth and they become progressively functional along the cochlea’s tonotopic gradient: transduction currents appear first at birth or postnatal day 0 (P0) in hair cells at the base of the cochlea, and from P2 in those at the apex (13, 14). Four salient aspects of MET have been observed to evolve with postnatal development. First, the peak transduction current increases. Second, the interval of hair-bundle displacements over which channels gate decreases. Next, channels open typically at smaller displacements with respect to the resting position of the hair bundle. Finally, adaptation becomes faster and more complete (13, 14).

Apparently unrelated to these changes throughout post-natal development is the dependence of fast adaptation on other experimental conditions. When measured at intermediate developmental stages, the speed of fast adaptation in a given cell decreases as a function of the magnitude of the imposed displacement, as can be seen from currents recorded in rat outer hair cells (Fig.5E in (14)) and in mouse inner hair cells (Fig.3C in (15)). In addition, hair cells with larger maximum MET currents display faster adaptation (Fig.2 in (9)). These results are not limited to mammalian hair cells and are not restricted to postnatal development: similar observations have been made in turtle hair cells (16, 17) and in embryonic chicken hair cells (12), suggesting a generic property of MET.

Similar trends have been observed in hair cells recovering from the severance of their tip links, a common effect of exposure to loud sounds (15, 18, 19). Tip links regenerate, and mechanosensitivity is fully recovered after 48 hours in mouse hair cells in culture (15, 19). During tip-link recovery over two days, the MET current undergoes changes that mirror those in developing hair cells over seven days: At first, the cell shows no mechanotransduction. After six hours, a small transduction current exists, but it displays little or no adaptation. As regeneration continues, the peak current increases, adaptation appears and then becomes progressively faster and more complete (15).

These results are difficult to explain within the standard theoretical framework of MET—the classical gating-spring (GS) model—which is fundamentally a single-channel model (2, 4). It is unclear, in particular, how fast adaptation could evolve during postnatal development or during tip-link regeneration, given that it is likely an intrinsic property of MET channels (14, 20, 21). One hypothesis is that a change in hair-cell Ca^2+^ buffering capacity due to a temporal gradient of PMCA2 (Ca^2+^ pump) expression, or perhaps a change in the subunit composition of the channel, could lead to the observed changes of the kinetics (13, 22, 23). For example, outer hair cells differentially express two members of the transmembrane channel-like (TMC) protein family during development (24–26). These proteins are believed to be pore-forming subunits of the MET channel and show different affinities to Ca^2+^, which could affect adaptation (22, 23). However, in apical hair cells in the mouse, TMC1 proteins only begin to localize to stereocilia tips at P6, whereas by that time the kinetics of adaptation have already matured (27). Moreover, changes in a protein expression in general cannot explain the corresponding phenomena in hair cells recovering from a loss of tip links, since tip-link regeneration and recovery of mechanotransduction do not require new protein synthesis (18). They also cannot explain why at intermediate stages of development adaptation is slower for larger imposed displacements (14, 15), and why it is faster for hair cells that display larger maximal currents (9).

Here, we propose a unifying explanation for the changes in the biophysical properties of the MET current during maturation of mechanotransduction, both in developing hair cells and during tip-link regeneration, as well as for the dependence of the kinetics of fast adaptation on the imposed hair-bundle displacement. We show that these phenomena can all be explained by the existence of two channel populations with different transduction properties. First, in a developing hair cells or one recovering after acoustic trauma, the relative proportion of these two populations varies with time, therefore changing the hair cell’s biophysical properties. Second, at a given time during development or tip-link regeneration, these populations are fixed but engage differently as a function of hair-bundle displacement, changing the average adaptation kinetics.

### THEORETICAL DESCRIPTION OF POSTNATAL MET MATURATION

In the experimental data obtained by Lelli *et al.* (13) from outer hair cells of the mouse cochlea, the number of functional MET channels increases during postnatal development, as indicated both by the increased current amplitude and uptake of the fluorescent dye FM1-43. More precisely, there is no transduction current before birth, and, by about P6 to P8, mechanotransduction reaches its adult characteristics. Similar observations have been made by Waguespack *et al.* (14) in outer hair cells of the rat cochlea from P0 to about P7.

It is not known precisely how the MET machinery is formed, nor is it known how many channels each tip link is connected to in the fully mature hair bundle. Here, in agreement with structural and functional data (28, 29) and as proposed in our recent model (30), we assume one channel per tip-link branch in mature hair cells, corresponding to two channels per tip link. We further hypothesize that, during the maturation process, individual MET channels connect to the tip links stochastically, populating the remaining available tip-link sites one by one with equal rates. As a result, at any time step during postnatal development, each tip link can find itself in one of the three following configurations: 1) the tip link is not connected to any channel, 2) the tip link is connected to a single channel as in the classical GS model, forming a ‘single-channel transduction unit’, 3) the tip link is connected to two channels as in the model of ref. (30), forming a ‘paired-channel transduction unit’. Following these rules, we illustrate in Fig. 1 a possible distribution of functional MET channels in a hair bundle at an intermediate stage of development.

**Figure 1:**
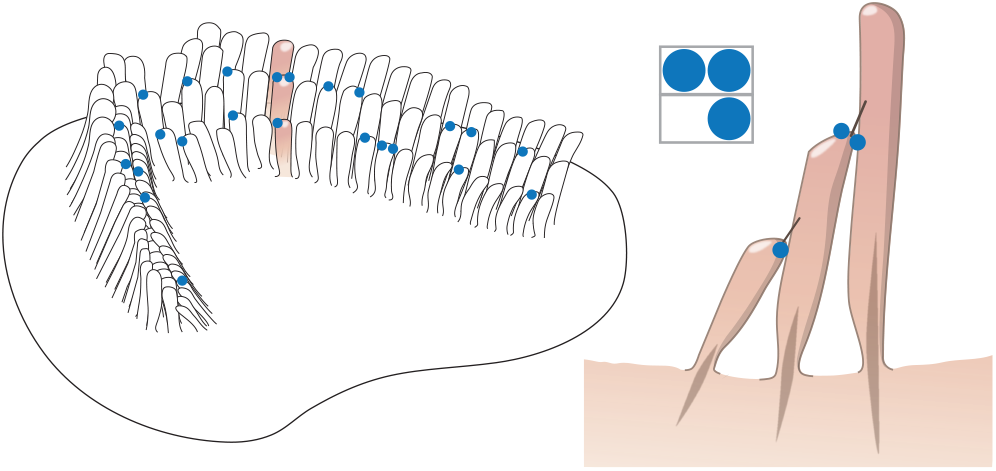
Possible distribution of MET channels in a hair bundle at an intermediate stage of development. Each blue dot represents a functional MET channel. Each tip link connects to either zero, one, or two channels.

### Population growth of MET channels

To distribute the channels across the hair bundle at any given time during postnatal development, one needs to specify how their number increases with the developmental stage of the cell. The simplest model is a linear increase, starting with zero channels at P0. Following this rule, the number of channels *n*_ch_ at postnatal day ζ (Pζ) reads:

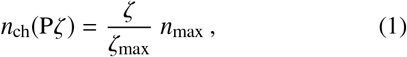

where *n*_max_ is the total number of MET channels present in the mature hair cell and ζ_max_ is the number of days required to complete the maturation process. In the experiments of Lelli *et al.* (13) and Waguespack *et al.* (14), however, the peak MET current increases sigmoidally during development. Assuming a constant single-channel conductance, a more realistic model is therefore a sigmoidal increase in the number of channels, as described by the logistic function:

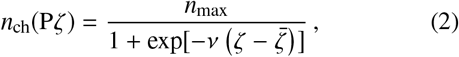

where *ν* and 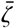 are fitting parameters. Fitting the curves reported in ref. (14) with a maturation time of seven days, we obtain 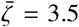 and *ν* = 1.26 per day. For a hair bundle with 50 tip links (4) and a maximum of two channels per tip link, we further take *n*_max_ = 100 channels. The data of ref. (13) for the mouse cochlea, however, point to a maturation time of six days. In that case, the same dependence applies with a simple rescaling of the phenomenological parameters, leading to 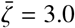 days and ν = 1.46 per day. In Table S1, we report the values given respectively by Eqs. 1 and 2 with *n*_max_ = 100 and both total maturation times of six and seven days, rounded up at each developmental stage to the closest integer number. In the following, we aim to compare our simulation results with the data of Lelli *et al.* (13) and Waguespack *et al.* (14). We therefore use the values reported here for the sigmoidal increase in the number of channels, with a maturation time of six and seven days, respectively.

### The most likely number of channel pairs

To simulate how mechanotransduction changes with hair-cell development, we need to specify how these MET channels distribute across the hair bundle’s transduction sites at each developmental stage. For a hair bundle comprising *n*_t_ tip links connected to a total of *n*_ch_ channels, with a maximum of two channels per tip link, the distribution of channels is set by the number *n*_p_ of paired-channel units and the following constraints:

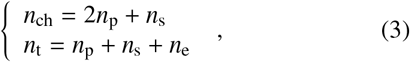

where *n*_s_ is the number of single-channel units and *n*_e_ that of tip links connecting to zero channels. Assuming that the channels are added one by one with equal probability to any of the remaining empty sites, the probability of having *n*_p_ formed pairs in the hair bundle reads:

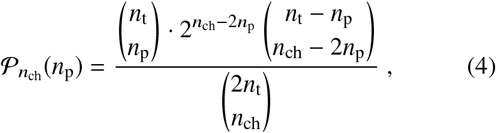

where the parentheses represent the binomial distribution (31). This equation states that once the *n*_p_ pairs have been distributed across the *n*_t_ tip links, *n*_ch_ − 2*n*_p_ single channels remain to be distributed across the *n*_t_ − *n*_p_ available tip links, with two choices per channel. The denominator corresponds to the total number of possible distributions of *n*_ch_ channels in 2*n*_t_ locations. This probability distribution is represented in Fig. 2 as a heat map for each value of the total number of channels *n*_ch_ between 0 and 100. In the following, we assume that the distribution of channels in the hair bundle corresponds to the configuration of maximum likelihood derived from Eq. 4. The white dots in Fig. 2 indicate the most likely state at each postnatal day from P0 to P6, assuming a sigmoidal growth of the number of channels in a hair bundle with 50 tip links that develops in six days. In Table S1, we report the values of the most likely number of channel pairs 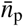, rounded up to the closest integer for every postnatal day from P0 to maturation (P6/P7), for both the linear and sigmoidal growths of the number of channels.

**Figure 2:**
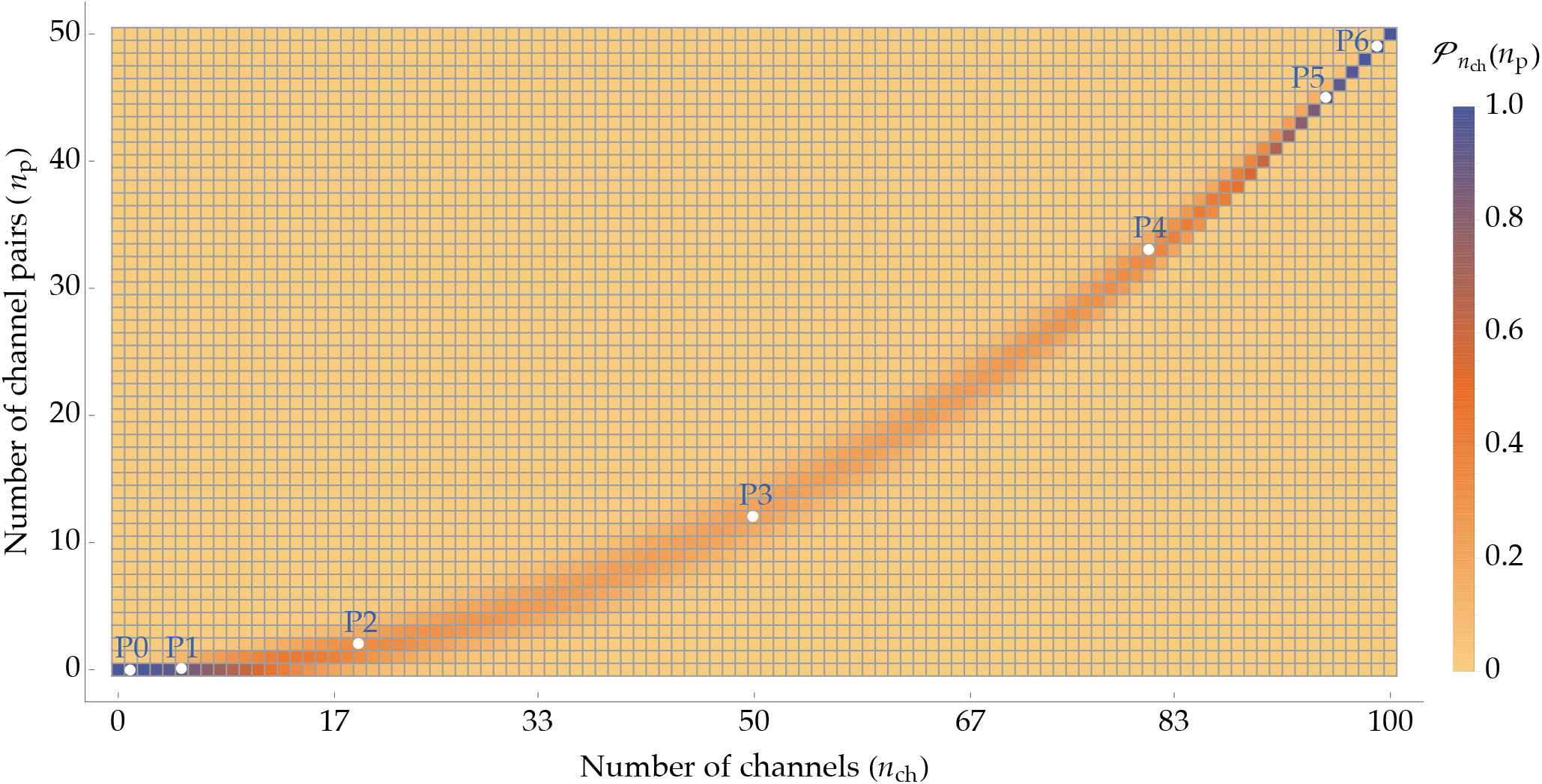
Heat map of the probability distribution of the number of channel pairs *n*_p_ for each value of the total number of channels *n*_ch_ as given by Eq. 4, in a hair bundle with 50 tip links (4). The white dots pinpoint the most likely numbers of channel pairs 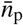 across development from P0 to P6, assuming a sigmoidal growth of the number of channels *n*_ch_ as given by Eq. 2, with *n*_max_ = 100 channels, 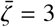, and *ν* = 1.46 per day.

### Open-probability curves

Having determined the most likely number of channel pairs 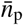 as a function of the developmental stage Pζ, we now study how the biophysical properties of the MET current change with development. In particular, we investigate the evolution of the open probability as a function of displacement at the level of the entire hair bundle and compare our model results with the experimental data of refs. (13) and (14).

The total response current of a hair cell is the sum of the currents through all open MET channels, which we assume to be identical. Single and paired units are expected to respond differently to hair-bundle displacements and are therefore described by different open-probability curves. This is because the open paired channels have an energetically favorable interaction through the lipid bilayer, a key contribution to the total energy balance of the system, which isolated channels lack. Therefore, channels in pairs open more readily than isolated ones. It follows that whenever the ratio of paired to single channels changes, the open-probability curve for the whole hair bundle changes accordingly.

To estimate the resulting open-probability curve, we rely on the finding that tip links within a hair bundle are mechanically coupled in parallel (32–36). We assume the same geometrical projection factor γ between the oblique orientation of each tip link and the axis of hair-bundle displacement, independently of the number of channels connected to that tip link (4). As a result, every unit is subjected to the same change of tip-link extension as a function of the hair-bundle displacement *X*. Therefore, the open-probability curve for the entire hair bundle at the developmental stage Pζ corresponds to the average contribution over all units present in the hair bundle. It reads:

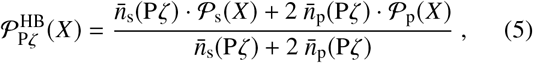

where 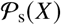 is the open-probability function of a single-channel unit, 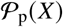 is that of a paired-channel unit, and 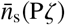 and 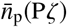 correspond to the most likely numbers of single- and paired-channel units at the developmental stage Pζ, respectively. The factor 2 accounts for the difference in channel numbers between single and paired units, and therefore for the difference in the contribution per unit to the total MET current, which is the experimentally measured quantity used to determine the open probability. We have discussed above how to determine 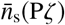 and 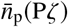. In the following paragraph, we specify our choice for 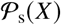 and 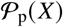

The open-probability curve 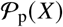 is derived from the paired-channel model introduced earlier, fully described in our previous work (30). Briefly, this model proposes that every tip link in a mature hair bundle is connected to two channels, as suggested by experiments (28, 37). Membrane deformations following opening and closing of either of the two channels couple their open probabilities, and the two channels gate cooperatively. This model reproduces the physiological behavior of the hair bundle without invoking an unrealistically large conformation change of the channel upon gating—the gating swing—as required when fitting with the classical GS model (4, 10, 38, 39). Instead, the large change in tip-link extension upon channel gating results here from the relative movement of the paired channels in the membrane. Since it includes membrane-mediated interactions between the paired channels, this model is also in agreement with the observation that the lipid bilayer plays a role in modulating the channels’ gating properties as well as in tuning fast and slow adaptation (10, 40).

This model depends on a list of parameters specified in Table S2. Among these parameters, a first category encompasses those describing the structural and mechanical properties of the hair bundle. These parameters appear in the model for single-channel units as well, as described further below. They are the tip-link stiffness *k*_t_, the gating swing δ, the projection factor γ, the difference in the energy of one channel between its open and closed conformations (or gating energy) *E*_g_, the total number of tip links *N*, and the total stiffness of the ensemble of stereociliary pivots *K*_sp_ (4). In addition, we must define parameters specific to the two-channel model, which describes channels mobile within the membrane and interacting via its deformations. Among these parameters are the length *l* of the branches of the tip-link fork, the stiffness *k*_a_ of the adaptation springs that anchor the channels to the actin network inside stereocilia, as well as a set of parameters describing the energy contribution of the stereociliary membrane, which depend on the state of the channel pair: open-open (OO), open-closed (OC), or closed-closed (CC). We refer the reader to ref. (30) for a full presentation. The parameter values reported in Table S2 have been chosen to correspond to those measured in a recent biophysical study of mammalian cochlear hair cells (41), where all the parameters relevant to the present work have been estimated from the same dataset. The resulting open-probability curve is displayed in Fig. 3 (red curve), with the origin of the *X* axis such that the hair bundle is at rest at *X* = 0 when no external force is applied.

**Figure 3:**
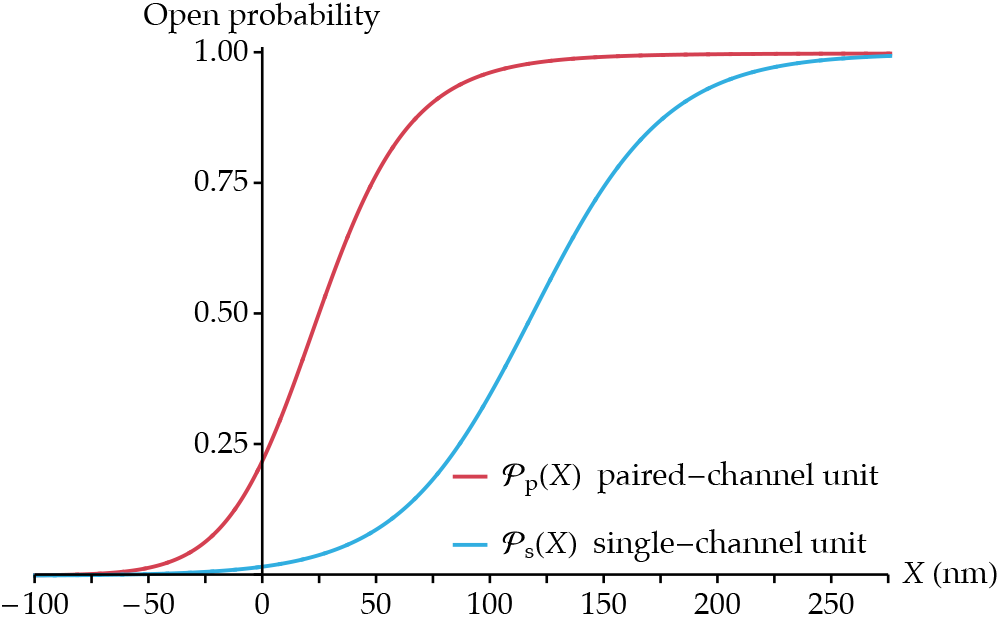
Curves of open-probability vs. hair-bundle displacement for the two types of units present in a developing hair bundle: paired-channel transduction units (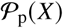, red curve) and single-channel transduction units (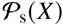, blue curve). The parameters that are shared by the two functions take the following values: *γ* = 0.1 and *k*_t_ = 0.7 mN⋅m^−1^ (41), δ = 2 nm (30), and *T* = 298 K. The parameters specific to the paired-channel model are set by our previous work (30) and are specified in Table S2. The origin of the *X* axis is such that the hair bundle is at rest at *X* = 0 in both models. This implies in particular *X*_1/2_ = 118 nm in Eq. 6 (see Eq. 1 in the Supporting Material).

We model the open-probability curve for isolated channels 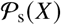 using the established GS model (4). Its analytic expression reads:

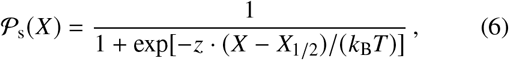

where *k*_B_*T* is the thermal energy at temperature *T*, *z* = *γk*_t_*δ* is the gating force, and *X*_1/2_ is the position along the *X*-axis for which half of the single channels are open (4). The value of *X*_1/2_ is set such that at rest the hair bundle sits at *X* = 0 when no external force is applied, similarly to the paired-channel model above. The resulting open-probability curve is shown in Fig. 3 (blue curve). We can see that, although the parameters common to both models have identical values, the open-probability curve from the two-channel model describes channels that gate at smaller displacements and is steeper than that from the classical GS model, corresponding to a higher sensitivity. These results are in agreement with studies performed on pairs of mechanosensitive channels of large conductance (MscL) in *Escherichia coli*, which have shown that closely apposed channels open at a smaller membrane tension than isolated ones and display steeper open probability curves, again corresponding to a higher sensitivity (42).

## RESULTS

### Leftward shift of the open-probability curve

The open-probability curve recorded from hair cells of the rat cochlea shifts toward smaller displacements during post-natal development (14). To compare our model results with these measurements, we plot in Fig. 4*A* the simulated open-probability vs. displacement curves across development as given by Eq. 5 in the case of a sigmoidal increase of the number of channels over six days (see Table S1). In Fig. 4*B*, we plot the derivatives of these open-probability curves to represent the hair-bundle sensitivity. A maximum of sensitivity corresponds to a maximum of the open-probability derivative, for which a maximum number of channels gate per unit hair-bundle displacement. At early developmental stages, most tip links connect to zero or one channel, and the open-probability curve resembles that of a single-channel unit (P1 curve here, to be compared with the 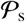 curve of Fig. 3). The sensitivity peaks for a displacement *X* around 125 nm at about 8.5 *µ*m^−1^, which corresponds to the maximal sensitivity of single channels. At late developmental stages, most tip links are connected to two channels, and the open-probability curves resemble that of a paired-channel unit (P6 curve here, to be compared with the 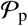 curve of Fig. 3). The sensitivity peaks for a displacement *X* around 25 nm where paired channels gate and is roughly 50% higher than that of single channels. At intermediate developmental stages, the open-probability curve appears as a weighted average of these two extremes and evolves with development by shifting toward smaller hair-bundle displacements as the number of single-channel units decreases and that of paired-channel units increases. Analogous results can be observed in the case of a linear growth in the number of channels. In Fig. S1, we report the evolution of the open probability curve across development (Fig. S1*A*) and its derivative (Fig. S1*B*) in a hair bundle reaching maturation linearly in six days (see Table S1).

**Figure 4:**
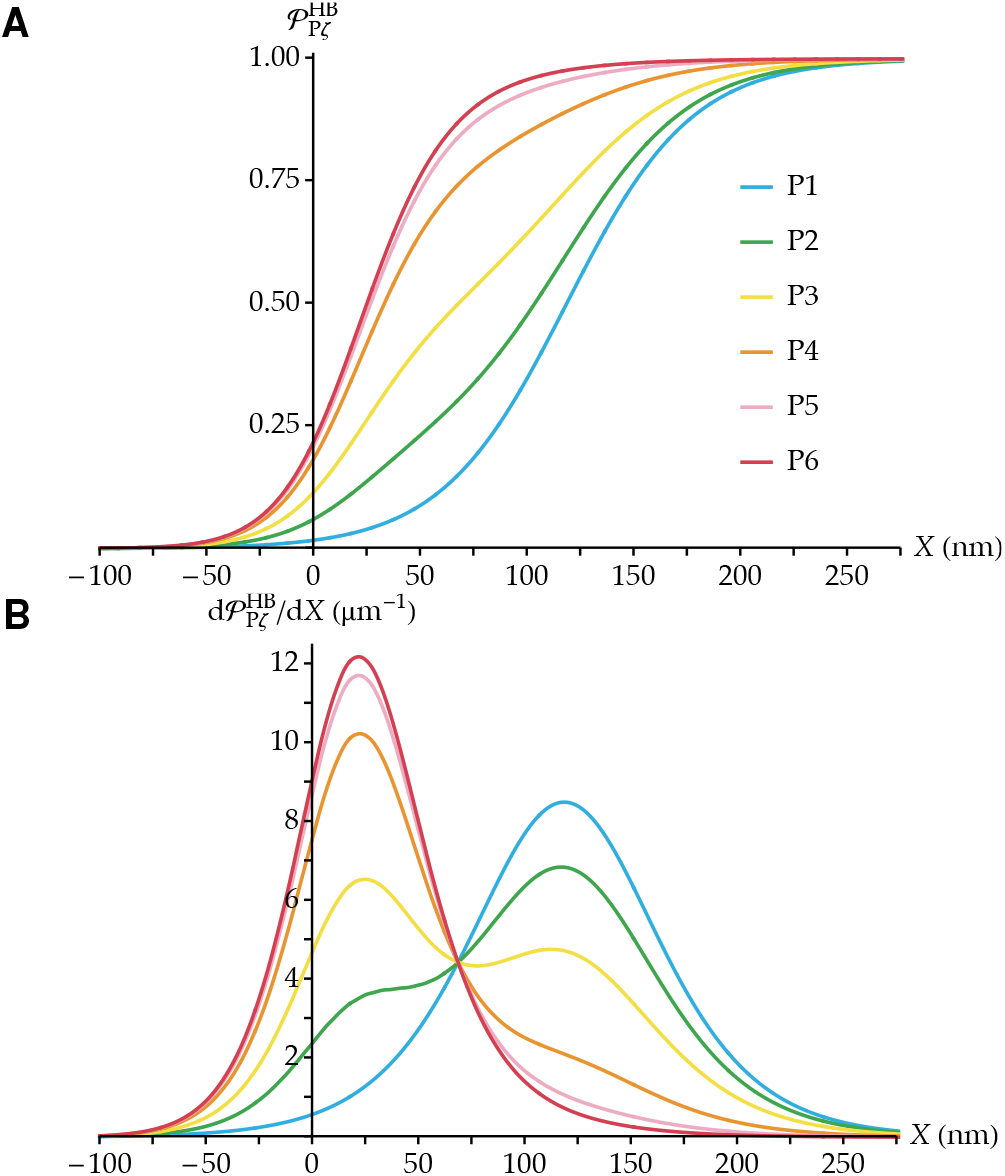
(A) Simulated maturation of the open-probability vs. displacement curve 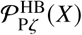 in a developing hair cell. We report the case of a sigmoidal growth of the number of MET channels over six days, as given in Table S1. The parameters that define the transduction units are the same as in Fig. 3. (B) Derivatives of the open probability curves of panel *A*, using the same color code. The open-probability curve shifts leftward with development; the hair cell becomes progressively more sensitive and its working range decreases accordingly.

Due to the presence of two different MET units that gate typically at different hair-bundle displacements and with two different sensitivities, the open-probability vs. displacement curves at intermediate stages of development present an asymmetric shape: compared to a standard sigmoid, they increase more rapidly at small displacements and then rise more gently to saturation, which resembles the shapes of the corresponding curves obtained experimentally (13). This phenomenon is comparable to what has been observed in voltage-dependent Ca^2+^ channels. In that case, data suggest the existence of two populations of ion channels as well, usually called ‘willing’ and ‘reluctant’ in that context (43). The asymmetric shape of the open probability curves is due to the fact that these two channel populations open at different voltages, similarly to what we propose here in terms of hair-bundle displacements.

### Decrease of the operating range

As outer hair cells mature, their operating range decreases and stabilizes by P6/P7 at a value that is roughly 50% that recorded at P0, both in rat and mouse cochleas (13, 14). This narrowing of the operating range is linked to an increase of the slope of the open-probability vs. displacement curve, reflecting an increased sensitivity of the hair cell to mechanical stimuli. To compare our simulation with these experimental data quantitatively, we choose as the operating range ∆*X*_op_ the difference in hair-bundle displacements corresponding respectively to 95% and 5% of open probability: 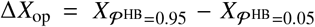. In Fig. 5, we plot this quantity at different developmental stages normalized by its maximum value. For a comparison, we report these results for both the linear and sigmoidal increase of the number of channels over six days (see Table S1).

**Figure 5:**
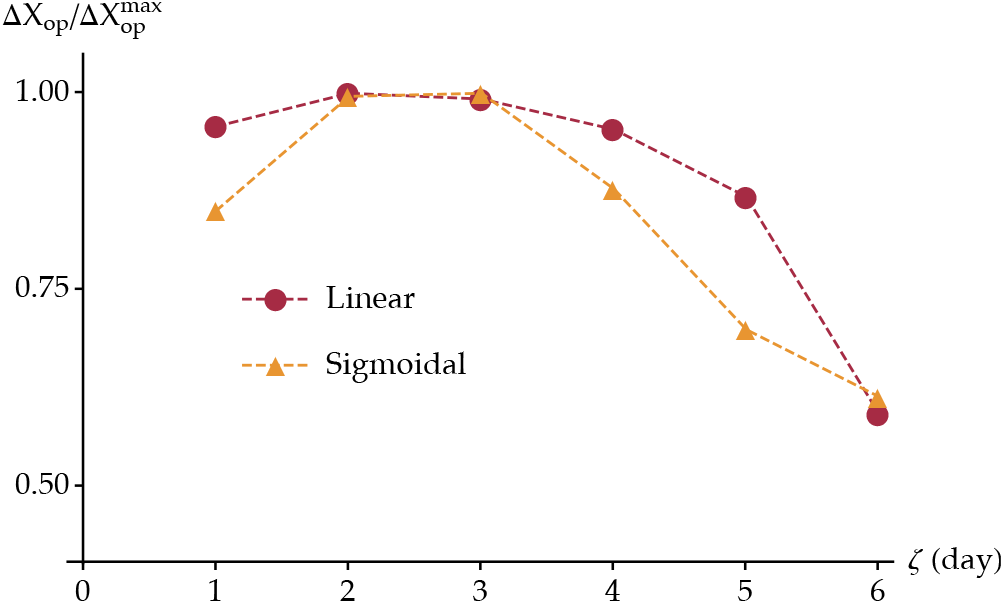
Simulated operating range normalized with respect to its maximum value and plotted between 0.50 and 1.00. Both cases of linear (red circles) and sigmoidal (orange triangles) growth functions of the number of channels over six days are reported.

### Maturation of adaptation

During development, adaptation becomes faster and its extent increases (13, 14). Its fast and slow components are characterized by two time constants, *τ*_f_ and *τ*_s_. They are obtained by fitting the response current *I*(*t*) of the hair cell as a function of the time *t* with the following expression:

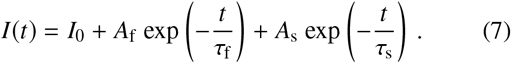

Here, *I*_0_ is a baseline current and *A*_f_ and *A*_s_ determine the contributions of fast and slow adaptation, respectively. Typical values of *τ*_f_ in mammalian hair cells range from 0.05 to 0.6 ms (8, 9), while *τ*_s_ ranges from 4 to 16 ms (8, 13, 14).

In rat and mouse hair cells, the time constants both decrease during early postnatal development (13, 14). In some cases, outer hair cells initially display mechanosensitivity but no form of adaptation (14, 44). Here, we propose that these results betray the existence of the two channel populations introduced above, which, in addition to having different open-probability vs. displacement curves, also have different adaptation time constants. Similarly as before, we propose that it is the change in their relative proportions that leads to the observed changes in the time constants of slow and fast adaptation at the level of the whole hair bundle.

Taking into account the existence of the two channel populations, the response current 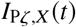 of the hair cell at a developmental stage Pζ and for an imposed step displacement *X* can be written:

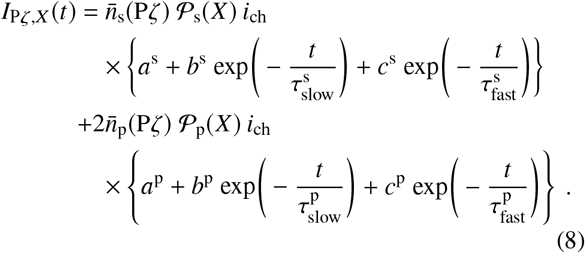

Here, 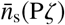 and 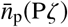 indicate the most likely numbers of single- and paired-channel units at Pζ, as discussed previously and reported in Table S1. The parameter *i*_ch_ is the current passing through one open MET channel, while 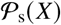 and 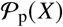 are the open probabilities of single and paired channels respectively, at the imposed displacement *X* and before adaptation takes place (see section “open-probability curves”). Together, the product 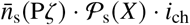 (respectively 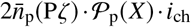) represents the total current passing through single channels (respectively paired channels) in the hair bundle before adaptation takes place. The dimensionless variables *b*^s(p)^ and *c*^s(p)^ weigh the relative contributions of the slow and fast components of adaptation for the single (s) or paired (p) channels, and *a*^s(p)^ = 1 − *b*^s(p)^ − *c*^s(p)^ gives the relative amplitude of the remaining currents after adaptation has taken place. The time constants 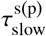 and 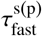 represent the characteristic times over which the channels undergo fast and slow adaptation, respectively, with specific values for single and paired channels. This equation states that the total decline in response current is the weighted sum of the contributions of the two types of MET units, each adapting at its own pace by the two independent processes of slow and fast adaptation.

To reduce the number of variables, we assume that the amplitudes of slow and fast adaptation are the same with *b*^s(p)^ = *c*^s(p)^. We then obtain:

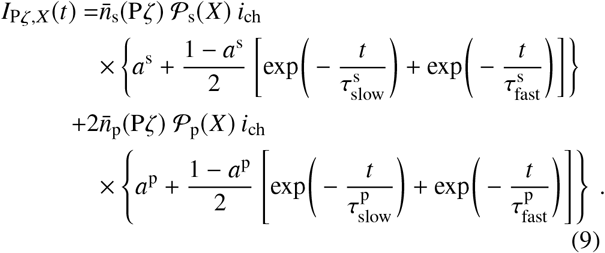

In the experimental data of refs. (13, 14), adaptive currents are measured by imposing step deflections onto the hair bundle such that the ensemble open probability is 50% at the onset of adaptation. To reproduce these results, we investigate the adaptation current for an imposed step displacement *X*_HB,50%_, at which the hair-bundle open probability 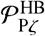 as given by Eq. 5 equals 50%, for each developmental stage Pζ. We set the parameters entering Eq. 9 by using the experimentally measured amplitudes of the MET currents before and after adaptation, both at early and late stages of hair-cell development. At P1, according to our model, most functional MET units contain only one channel (13, 14). Therefore, the currents at the onset of adaptation and after adaptation has taken place read, respectively:

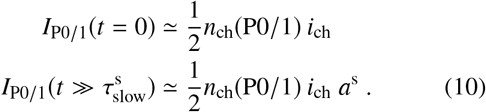

These expressions allow us to express the extent of adaptation, defined at the developmental stage Pζ as 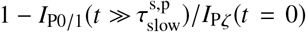 (13, 14). At P0/1, this quantity is simply (1 − *a*^s^). In addition, the time constants 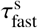 and 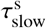 are taken directly from the time traces of the currents recorded at P1. The same reasoning leads to the corresponding quantities at the end of maturation, at P6 or P7 depending on which data set (13, 14) we consider:

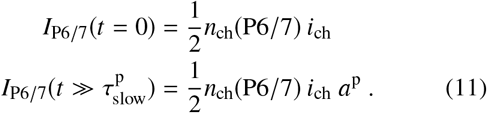

Similarly, the extent of adaptation at P6/7 is (1 − *a*^p^) and the time constants 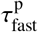 and 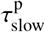 are estimated from the time traces recorded in mature hair cells (P6/P7).

In the basal region of the mouse cochlea, the extent of adaptation ranges from 80% at P0 to 95% at P6 (13), setting *a*^s^ = 0.20 and *a*^p^ = 0.05. In addition, we obtain 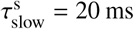 and 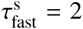 as the adaptation time constants measured at P1, and 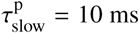 and 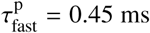 at P6 (13). We display in Fig. 6*A* the resulting simulated currents as given by Eq. 9, from P1 to P6. Each current is normalized with respect to its value with half of the channels open in the fully developed hair cell *I*_P6_(0) = *n*_ch_(P6)/2) × *i*_ch_, corresponding to the maximal current of the hair cell at the imposed displacement *X*_HB,50%_. We can see an inflection in the response-current curves after P3, where the double exponential becomes clearly visible, corresponding to two distinct kinetics of the overall fast and slow adaptation components at the level of the whole hair bundle.

**Figure 6:**
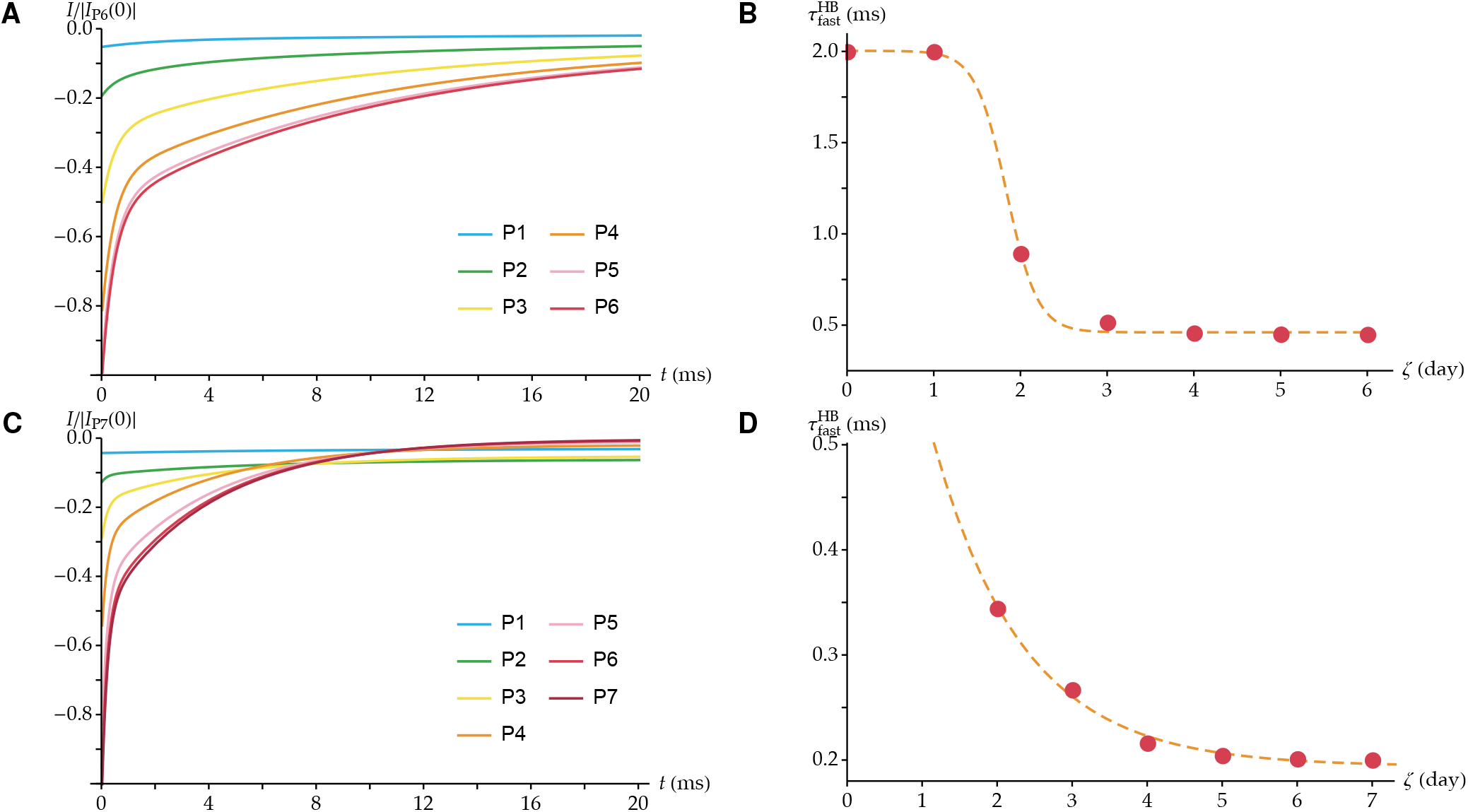
Normalized response currents (*A,C*) and time constants of fast adaptation for the whole hair bundle (*B,D*), across simulated postnatal development. (*A*) The response currents are generated using Eq. 9, with a sigmoidal growth of the number of channels over six days (see Table S1) and the following parameter values: 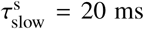, 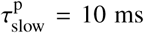, 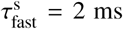, 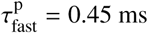, *a*^s^ = 0.20, and *a*^p^ = 0.05. For each curve, *X* is computed such that 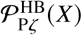 is equal to 0.5, as given by Eq. 5. (***B***)The time constant of fast adaptation at the level of the whole hair bundle 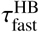 is reported as a function of the postnatal day Pζ, obtained by fitting the curves of panel *A* with Eq. 7. These results aim at reproducing the data in Lelli *et al.* from the mouse cochlea (13). (*C*) and (*D*) A similar procedure is followed to reproduce the data in Waguespack *et al.* from the rat cochlea (14). (***C***) The response currents are generated using Eq. 12 with a sigmoidal growth of the number of channels over seven days (see Table S1) and the following parameter values: 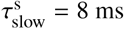, 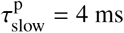, 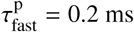, *a*^s^ = 0.7, and *a*^p^ = 0. In panels *B* and *D*, the results are displayed together with their best fits to a sigmoidal and an exponential functions, respectively, as done in refs. (13, 14) (dashed lines).

The simulated curves display a large variation in their apparent relaxation times. To quantify this variability, we fit these curves with a double exponential as in Eq. 7, with adaptation time constants 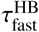 and 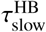 at the level of the whole hair bundle, as if the simulated curves were experimental data. We report in Fig. 6*B* the values of 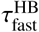 obtained across development, from P1 to P6. In addition, at P0, we have a single isolated channel in the simulated hair bundle, such that 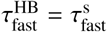. We can see that 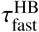 decreases with development, in agreement with the experimental findings (13). A similar global trend holds true for slow adaptation (see Fig. S2A). Overall, these results are in agreement with the experimental observations of ref. (13).

The same procedure can be applied to simulate adaptation as observed in the rat cochlea (14). Some of these experiments show an absence of fast adaptation at early stages of development. Within our framework, this suggests that isolated channels might only be capable of slow adaptation in that case. In contrast, the paired channels would still display both components of adaptation, as observed in the fully developed hair cells. Under these hypotheses, Eq. 9 is replaced by:

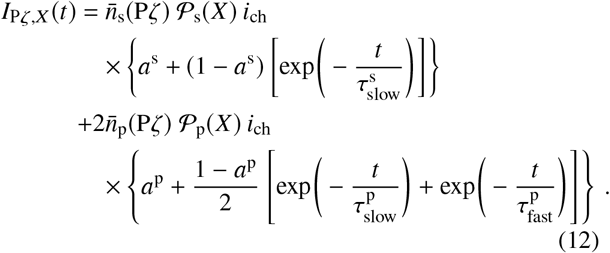

Experimentally, the characteristics of adaptation were only measured in the basal part of the rat cochlea for the cells that displayed fast adaptation at P1. Assuming that slow adaptation remains the same in all basal cells at that developmental stage, we can use these quantifications to simulate the cells that do not display fast adaptation at P1. These measurements indicate that the extent of adaptation is around 30% at P1 and reaches 100% by P3/P4 (14). This sets *a*^s^ = 0.7 and *a*^p^ = 0. In addition, the fitted kinetics of fast and slow adaptation at P1 and P7 give 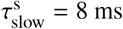, 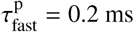, and 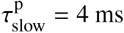.

With these parameters, we display in Fig. 6*C* the normalized adaptation currents at different developmental stages Pζ. In Fig. 6*D*, we report the corresponding fitted values of 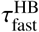 at each developmental stage, from P2 to P7. At P1, there are no paired channel units in the model and fast adaptation is absent. At later developmental stages, 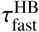 is finite. It then decreases with development and reaches about 0.2 ms at maturation. These results are in agreement with the trend observed in refs. (14, 44), where the onset of adaptation has been observed to lag that of mechanotransduction. Similarly to Fig. S2a, we report in Fig. S2b the corresponding decrease in 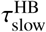 as a function of Pζ.

Data from Lelli *et al.* (13) suggest that, at P1, MET channels in mouse hair cells display fast adaptation, whereas Marcotti *et al.* observed hair cells incapable of adaptation at P2 in the same animal model (Fig. 7 in (44)). Waguespack *et al.* observed no fast adaptation in a fraction of rat hair cells at early postnatal developmental stages (14). In our analysis, we used single channels with or without fast adaptation to reproduce those experimental data, and we showed that the model accommodates either type of behavior. These apparent discrepancies within recorded data can be resolved assuming that fast adaptation at early stages of postnatal development results from paired-channel units already present in the hair bundle. At P1, rat basal hair cells in (14) display a peak current of about 100 pA at a holding potential of −80 mV. This is approximately one third of that recorded in mice of the same age at a holding potential of −64 mV (13), suggesting that the observed difference in the adaptation properties could be the result of different developmental stages at birth. Small currents (~ 150 pA at −84 mV) have also been recorded at P2 in mouse preparations by Marcotti *et al.* (44), where hair cells displayed no adaptation.

**Figure 7:**
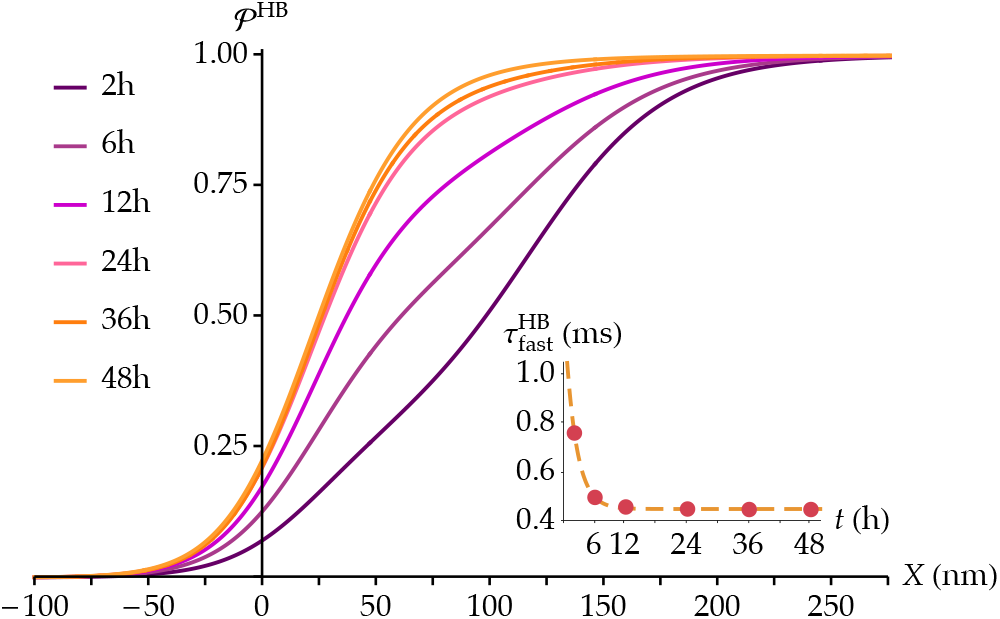
Evolution of the ensemble open-probability function in a simulated hair cell with 50 broken tip links after 2, 6, 12, 24, 36, and 48 hours of recovery (main), and corresponding evolution of 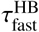 (inset). The curves are derived from Eqs. 5 and 9 with the number of channels from Eq. 14 and the following parameters: 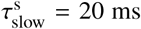, 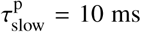, 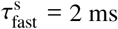, 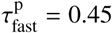, *a*^s^ = 0.20, and *a*^p^ = 0.05. The factors 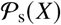 and 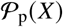 are calculated with an imposed displacement *X*_HB,50%_ such that 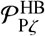 equals 50%.

In agreement with this interpretation, we show in Fig. S3 that a similar trend in the maturation of the time constant of fast adaptation as the one observed in (13) can be generated assuming that single-channel units are not capable of fast adaptation but that MET development starts before birth with paired channels already present at P1. In this figure, we represent 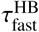 as a function of Pζ using the same parameter values as in Fig. 6D, but with a channel population growth over a new total of ten days and starting three days before birth to end at P7 as before. The hair cell displays a finite value of 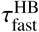 already at P0, as in (13) and our earlier Fig. 6B, despite the fact that single-channel units in this case do not fast adapt.

Our model also explains why it is possible to observe fast adaptation in some hair cells but not in others obtained from the same animal. At early developmental stages, the number of channel pairs *n*_p_ can vary from cell to cell, the actual values being taken from the probability distribution given by Eq. 4 and illustrated in Fig. 2. This means that, at early developmental stages, some hair bundles can contain paired-channel units, therefore displaying a certain amount of fast adaptation, while others do not.

### Tip-link regeneration

One of the effects of loud sounds on hair cells is the disruption of tip links. This effect is temporary, however, because tip links can regenerate (15, 18, 19, 45). In mouse hair cells in culture, this process takes about 48 hours (15), during which the biophysical changes of the MET current recapitulate those observed in postnatal development across seven days: the peak current becomes larger, adaptation becomes faster and more complete, and the slope of the open probability-displacement curve increases (15, 18, 19).

To restore hair-cell mechanosensitivity, new tip links must establish connections with MET channels. We simulate this process by assuming that each of the tip link’s two branches has a given probability per unit time 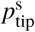 to connect to a MET channel, and a probability per unit time 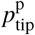 that the second branch connects to another MET channel, given that the first branch has already been linked. The evolution of the number of paired- and single-channel units is determined by the following equations:

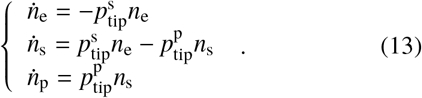

Here, 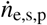 represent the time derivatives d*n*_e,s,p_/d*t* respectively, where *t* is the time passed after tip-link damage, on the order of several hours. For simplicity, we consider the case 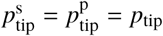. Eq.13 is then solved by:

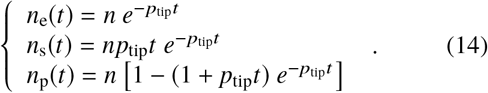

The evolution of these quantities allows us to simulate the change in the open probability curve in a hair cell recovering from the loss of its tip links using Eq. 5, replacing 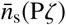 and 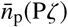 by *n*_s_(*t*) and *n*_p_(*t*) as given here, respectively. We further simulate the evolution of fast adaptation using Eq. 9, as described above and with the parameters derived from the experiments of ref. (13) on mouse cochlear hair cells. The value of *p*_tip_ is chosen so that the hair bundle has fully recovered by *t* = 48 h. Since the above model reaches *n*_p_ = 50 only asymptotically at *t* = ∞, we consider that full recovery is reached when *n*_p_ (*t*) > 49.5, such that the closest integer value is 50. Setting that this value is reached at 48 h leads to *p*_tip_ ≃ 0.14. In Fig. 7, we show the resulting open-probability curves at 2, 6, 12, 24, 36, and 48 hours after tip-link severing. In the inset, we show the change in the fast-adaptation time constant at the level of the entire hair bundle, following the same fitting procedure as the one used in the previous section. The results agree with those observed experimentally in the mouse cochlea (15, 19): the slope of the open probability curve increases during tip-link regeneration, similarly to the increase during postnatal development (see Fig. 4), while adaptation becomes faster, again resembling the change in the kinetics during development (see Fig. 6D).

### Adaptation kinetics as a function of the transduction-current amplitude

At intermediate developmental stages, where both types of units are present, we expect adaptation to become slower with larger imposed displacements. This is because larger displacements open a larger proportion of channels in single-channel units, which adapt more slowly. Such a behavior has been observed experimentally in cochlear hair cells of mice (9, 13, 15) and rats (Fig.5E in ref. (14)) several days after birth. In fully mature hair cells, where we predict that all tip links are connected to the same type of units, this dependency should vanish. In mature hair cells from the mouse utricle and from the frog sacculus, the time constant of fast adaptation is indeed independent of displacement (46, 47). Moreover, according to our model, hair cells with a greater total number of channels have also a greater fraction of them in pairs. Hence, adaptation is expected to be faster for cells that exhibit a larger maximum transduction current. Such a behavior has been reported in Fig. 2b of ref. (9).

In Fig. 8, we investigate the kinetics of fast adaptation as a function of the transduction-current amplitude using Eq. 9, at different developmental stages. In Fig. 8A, we plot the time constant of fast adaptation for the whole hair bundle 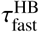 as a function of *I*_Pζ,X_ (0)/*I*_max_ for each developmental stage Pζ between P2 and P7 and imposed displacement *X*. Here, *I*_Pζ,X_(0) is the negative peak current for a given displacement *X* at the onset of adaptation (t=0) in a hair cell at the developmental stage Pζ, and *I*_max_ = *n*_ch_(P7)× *i*_ch_ is the maximal value of the transduction current at maturation. Note that we exclude from this analysis the developmental stages P0 and P1, for which paired channels are not always present. At each developmental stage Pζ, each data point corresponds to a different value of the overall open probability 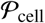 at *t* = 0, from 10% to 100%, in steps of 10%. We can see that the amplitude of the imposed hair-bundle displacement, directly related to the value of the current *I*_Pζ,X_(0), has a strong influence on the adaptation kinetics at early developmental stages of the cell, and progressively less influence at later developmental stages. At P7, as only paired channels are present, this influence completely disappears. At P2, however, the effective time constant more than doubles between the two extreme values. This trend is in agreement with the results reported in mouse inner hair cells (15) as well as in rat outer hair cells (14).

**Figure 8:**
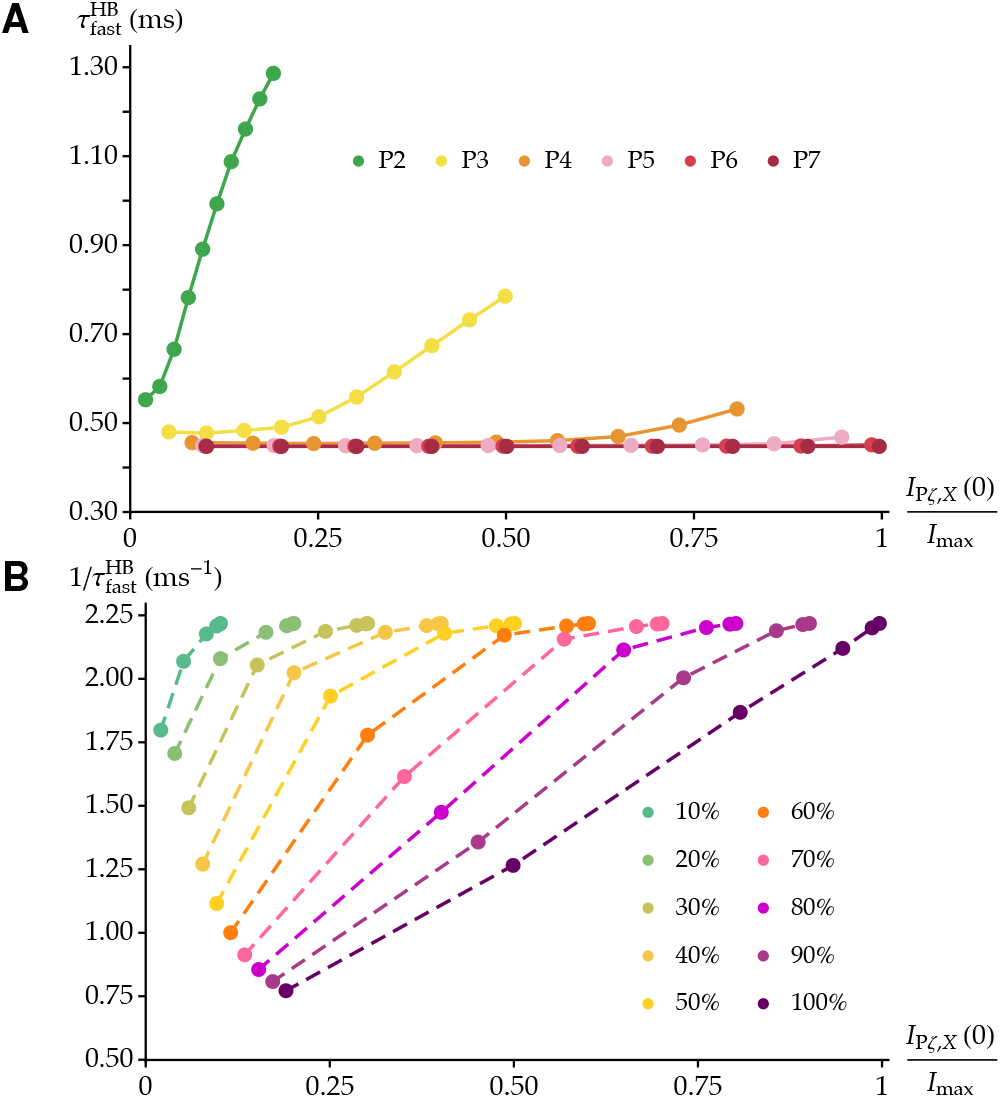
Adaptation kinetics as a function of the normalized peak transduction current at each developmental stage and imposed displacement. (A) The time constant of fast adaptation for the whole hair bundle 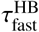 is plotted as a function of the normalized current at *t* = 0 as the displacement *X* is varied, at each developmental stage Pζ from P2 to P7. In each series, each data point corresponds to a different value of the overall open probability before adaptation 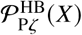, as given by Eq. 5, from 10% to 100% in steps of 10%. (B) The rate of fast adaptation 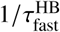 is plotted as a function of the same quantity as in (A), this time gathered per equal values of 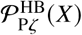. The plots are generated with the parameters characterizing the data in Lelli *et al.* from the mouse cochlea (13), as in Fig. 6A,B. Specifically, we assume a sigmoidal growth in the number of channels over six days as reported in Table S1, and the other parameter values are: 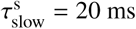, 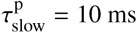, 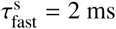, 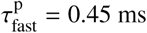, 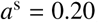, and *a*^p^ = 0.05.

To compare with other experimental data (9), we plot in Fig. 8B the rate of fast adaptation 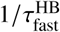 as a function of the same quantity as in Fig. 8A. We group these values per equal open probability, and we can see that, at fixed open probability, the rate of adaptation increases with the peak current across development, in agreement with the experimental results of ref. (9). When using parameters derived from the data in Waguespack *et al.* (14) and Eq. 12, this analysis yields similar results (see Fig. S4).

## DISCUSSION

The maturation of mechanotransduction in cochlear hair cells across several days during postnatal development, as well as across several hours during tip-link regeneration, is characterized by a number of hitherto unexplained variations in the MET currents’ biophysical properties. Instead of relying on the differential expression of molecularly distinct MET channels, we have explained these changes as the result of a random connection of tip links to a population of MET channels with identical and constant biophysical properties. In this framework, the maturation of the MET current’s biophysical properties is the result of a change in the relative numbers of transduction units that contain either a single or a pair of channels. Although composed of the same kind of channels, these two types of units behave differently because channels in pairs interact with one another via the membrane bilayer, which substantially affects their open probability (30). Single channels display relatively broad open-probability curves, centered at relatively large hair-bundle displacements, whereas paired channels display steeper open-probability curves, centered at smaller displacements.

Using this framework, we have reproduced the following aspects of hair-cell postnatal development: First, as development proceeds, hair cells become more sensitive in that the channels gate at smaller hair-bundle displacements and over a narrower range of displacements. Second, at intermediate developmental stages, the transduction-current vs. displacement curves have an asymmetric shape, increasing more rapidly in the first half of their gating range. Third, the time constants of fast and slow adaptation decrease during development. Fourth, at intermediate developmental stages, the time constant of fast adaptation depends on the amplitude of the imposed hair-bundle displacement. In addition, we showed that we could reproduce the similar biophysical changes observed during tip-link regeneration using the same model.

To reproduce the change in the measured fast-adaptation kinetics, we hypothesized that the adaptation time constants for single and paired channels are different. Published ensemble-averaged traces based on single-channel recordings suggest that single channels do not always adapt (Fig.5 in (48)). More specifically, ensemble-averaged current traces in a hair cell from the apex of the mouse cochlea at P2 show no adaptation. As these recordings represent the activity of a MET unit whose conductance is at the lower end of the observed range, they are most likely due to the activity of a single channel. In contrast, prominent adaptation has been observed in ensemble-averaged traces from relatively more mature basal hair cell at P2 and apical hair cell at P4 (29). The associated conductance values sit in the middle or at upper end of the measured range, respectively, which, within our framework, indicates that they reflect the concerted opening of more than one channel per MET unit, as suggested by the authors of that study. Another series of similar recordings from mouse outer hair cells at P4 and P6 showed current traces with several peaks in the amplitude histograms, interpreted as the concerted openings of different numbers of MET channels (29). Those records showed a prominent adaptation as well.

To see if paired channels could produce fast-adaptation kinetics consistent with those observed in mature hair cells, we attempt to interpret the fast-adaptation time constant within the framework of our two-channel model. Within the reaction-rate theory, the kinetics of fast adaptation is limited by a transition state with an associated activation energy *E*_a_. In this context, the rate of fast adaptation *k*_fast_ is given by the empirical law of Arrhenius:

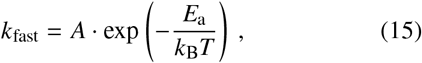

where *k*_B_ is the Boltzmann constant, *T* is the absolute temperature, and *A* is the so-called attempt frequency. In the following, we start by estimating the attempt frequency and then use the measured adaptation time constants to derive estimates for the activation energy, which we can interpret in the context of our two-channel model.

To estimate the attempt frequency *A*, we relate it to the rate of permeation of Ca^2+^ ions through an open MET channel, which in turn can be approximated by:

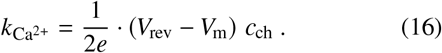

Here, *e* is the elementary charge, *V*_rev_ is the reversal potential of Ca^2+^, *V*_m_ is the membrane potential of the cell and *c*_ch_ is the conductance of the MET channel. At an extracellular Ca^2+^ concentration of 1.5 mM, 15% of the total current is carried by Ca^2+^ (49). Therefore, *c*_ch_ should be 15% of 50 pS, which is the conductance of a TMC1 pore subunit (29). Together with *V*_rev_ ≃ 75 mV and *V*_m_ varying in a range from −60 mV to −85 mV (13, 14, 44), one finally gets an attempt frequency per channel *A*_s_ ≃ *k*_Ca_2+ ranging from 3 − 4 ⋅ 10^6^ s^−1^. Using the membrane potential *V*_m_ = −64 mV as in (13), we obtain an attempt frequency for single channels *A*_s_ ≃ 3.3 ⋅ 10^6^ s^−1^. For paired channels, the rate of Ca^2+^ entry needs to be doubled, which leads to *A*_p_ ≃ 6.5 ⋅ 10^6^ s^−1^. With a hyperpolarized membrane potential *V*_m_ = −80 mV (14), the attempt frequencies read *A*_s_ ≃ 3.6 ⋅ 10^6^ s^−1^ for single channels and *A*_p_ ≃ 7.3 ⋅ 10^6^ s^−1^ for paired channels.

We now evaluate the activation energy *E*_a_ using the estimations above and the measured time constants. In basal outer hair cells of the mouse cochlea at P7, the fast-adaptation time constant is 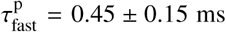 (13). Together with our estimate of the attempt frequency *A*_p_ ≃ 6.5 ⋅ 10^6^ s^−1^ at *V*_m_ = −64 mV, we obtain an activation energy *E*_a_ ~ 8.0 *k*_B_*T*. For *V*_m_ = −80 mV the result is very similar, around 8.1 *k*_B_*T*. Considering that the work done by the binding of one calcium ion is *W*_Ca_^2+^ ≃ 7 *k*_B_*T* (50), one obtains an energy barrier in the absence of calcium of about 15 *k*_B_*T*.

To interpret this result in the context of our two-channel model (30), we now estimate the energy required to close a pair of ion channels in the open-open configuration. Due to the membrane-mediated interaction, the two channels are in contact with each other before adaptation. This allows us to compute the energy of the system as the sum of the following contributions: The energy difference of one adaptation spring between the closed and open states of the channel is [(1/2)*k*_a_(*e*_a,max_)^2^ − (1/2)*k*_a_ (*e*_a,max_ − δ)^2^] in favor of the open state, where *e*_a,max_ is the extension of the adaptation springs when the channels touch in the closed-closed (CC) configuration, δ is the variation of the channel lateral width upon gating, and *k*_a_ is the stiffness of the adaptation springs. With the default parameters reported in Table S2, we have *e*_a,max_ ≃ 12 nm (see ref. (30)), which leads to an energy difference of 5.4 *k*_B_*T* in favor of the open state. With the contribution from the channel gating energy *E*_g_ = 9 *k*_B_*T*, we obtain a total contribution of 3.6 *k*_B_*T* per channel in favor of the closed state, or 7.2 *k*_B_*T* for the pair. Finally, the contribution from the membrane elastic energies is a difference of 22.5 *k*_B_*T* in favor of the open-open (OO) configuration vs. the CC one. Altogether, the OO state is therefore favored by about 15.3 *k*_B_*T* over the CC state at the point where the paired channels are in contact, a value that agrees with the one estimated from the measurements of the adaptation rates above. After the channels have closed, the adaptation springs pull them apart, stabilizing the CC configuration. It is therefore reasonable to consider the CC state with the two channels in contact as the activation state for channel closing. The fact that both the direct measurements of fast-adaptation time constants and the aforementioned molecular schema derived from our two-channel model yield similar estimates reinforces this interpretation and supports the conclusion that paired channels can be at the origin of the measured kinetics of fast adaptation in mature hair cells.

In the model developed in ref. (30) and used here, energy is required not only to open the paired channels but also to close them. This is because, when the channels are in contact, the OO state is stable independently of tip-link tension, due to the favorable membrane-mediated interaction. This result has two implications: First, mechanotransduction in a fully developed hair bundle requires an external energy source. Second, mechanotransduction and fast adaptation are both active processes, relying on the same external energy source, potentially the out-of-equilibrium gradient of Ca^2+^ concentration across the cell membrane (50). Thanks to such an active process, the paired channels would close and the adaptation springs would separate them, in turn affecting tip-link tension and ultimately performing mechanical work on the hair bundle on a cycle-by-cycle basis. Other, passive properties related to fast adaptation, such as tip-link viscoelasticity (51, 52), could coexist with this active process. However, they cannot replace it. In contrast, single channels do not require energy to close, but can do so passively whenever tip-link tension is reduced. This allows the transduction current to passively follow the external stimulus on a cycle-by-cycle basis. Physiologically, this property could be beneficial for hair-cell development, which requires some resting MET current to progress normally (53). As the MET machinery fully matures and single-channel units are replaced by paired-channel ones, the active process would progressively come into play, and mechanotransduction and amplification become intimately linked.

## AUTHOR CONTRIBUTIONS

F.G. designed and developed the theory, performed the calculations and the simulations, produced the figures, interpreted the results, and wrote the manuscript. T.R. supervised the development of the theory, interpreted the results, and wrote the manuscript. A.S.K. conceived the proposed mechanism, designed and supervised the project, interpreted the results, and wrote the manuscript. All authors contributed extensively to this work.

## ACKNOWLEDGMENTS

Work on this project in the A.S.K. lab is funded by The Royal Society (*RG*140650), the Wellcome Trust (108034 *Z* 15 *Z* and 214234 *Z* 18 *Z*) and the Imperial College Network of Excellence Award.

## Supporting Information

### LINEAR AND SIGMOIDAL GROWTHS IN THE NUMBER OF CHANNELS

**Table 1:**
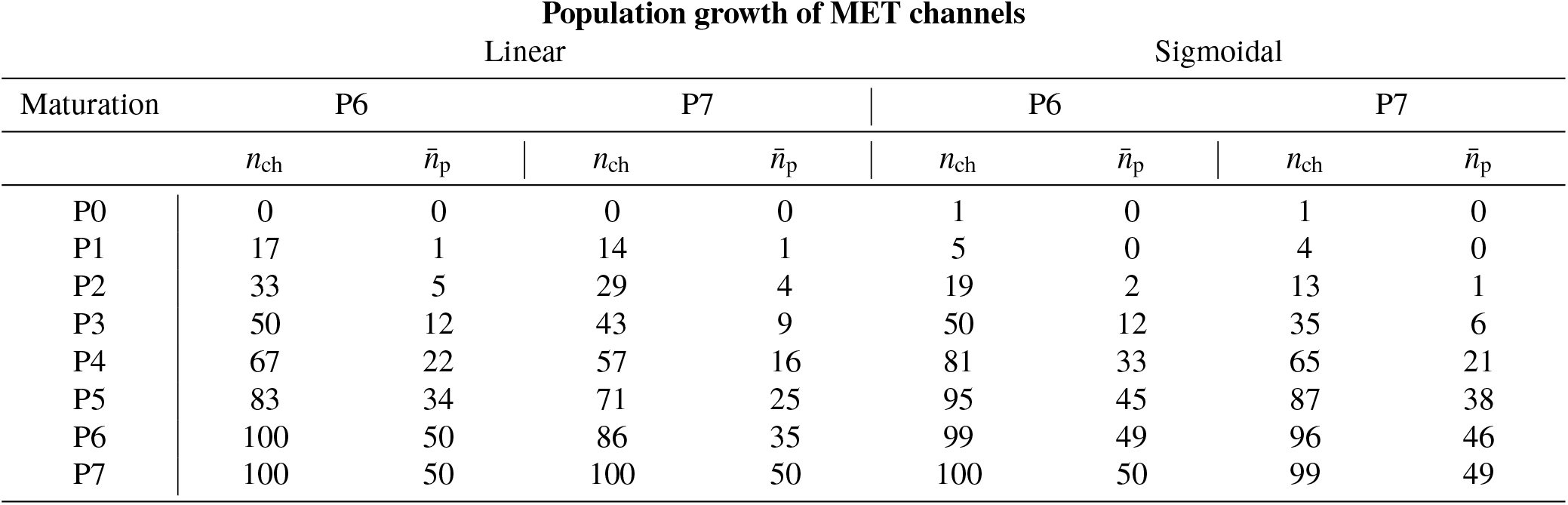
Number of channels *n*_ch_ and associated most-likely number of channel pairs 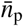 in a hair bundle with 50 tip links, corresponding to *n*_max_ = 100 channels, as a function of the developmental stage Pζ. Two different types of growths of the total number of channels are reported: a linear and a sigmoidal growths, as dictated by Eqs.1 and 2 of the main text, respectively. For each of these growth models, we simulate the development over a total maturation time ζ_max_ of either six or seven days, to account for the variations observed experimentally. For the sigmoidal growth, we have 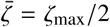 and ν = 1.26 or 1.46 per day, respectively for these two cases. For each of the four obtained population growths, the most likely number of channel pairs 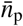 is given by the maximization of Eq. 4 of the main text at fixed *n*_ch_ and *n*_t_, rounded up to the closest integer value.

### PARAMETERS OF THE MODEL

**Table 2:** List of parameters

| Parameters characterizing the hair bundle and the mechanotransduction units |  |  |  |  |
| --- | --- | --- | --- | --- |
| Par. | Description |  | Default value | Unit |
| $k_t$ | Tip link's stiffness | Tobin 2018 (1) | 0.7 | mN·m<sup>-1</sup> |
| $\delta$ | Channel's steric change upon gating | Ursell 2007 (2) | 2 | nm |
| $\gamma$ | Geometrical projection factor | Tobin 2018 (1) | 0.1 | adim. |
| $E_g$ | Channel's gating energy | Corey 1983 (3); Hudspeth 1992 (4); Ricci 2006 (5) | 9 | $k_B T$ |
| $N$ | Number of tip links | Howard 1988 (6); Tobin 2018 (1) | 50 | adim. |
| $K_{sp}$ | Combined stiffness of the stereociliary pivots | Tobin 2018 (1) | 2.1 | mN·m<sup>-1</sup> |
| $l$ | Length of the tip link's branch | Kachar 2000 (7), Araya-Secchi 2016 (8) | 13 | nm |
| $k_a$ | Adaptation spring's stiffness | Powers 2012 (9) | 1 | mN·m<sup>-1</sup> |
| $\alpha$ | Angle of the adaptation springs w.r.t the horizontal | Kachar 2000 (7); Powers 2012 (9) | 5 | degrees |
| $a_{adapt}$ | Value of $a$ with relaxed adaptation springs (CC config.) | Kachar 2000 (7) | 13 | nm |
| $\rho$ | Radius of the closed channel | Ursell 2007 (2) | 2.5 | nm |
| $a_{min}$ | Minimum value of $a$ | Ursell 2007 (2) | 1.25 | nm |
| $X_{sp}$ | Resting position of the pivots | Tobin 2018 (1) | 21 | nm |
| $X_{1/2}$ | Value of the hair-bundle displacement $X$ such that $\mathcal{P}_s(X) = 0.5$ | Howard 1988 (6) | 118 | nm |

| Parameters characterizing the elastic membrane potentials |  |  |  |  |
| --- | --- | --- | --- | --- |
| Par. | Description |  | Default value | Unit |
| $a_{cross,CC}$ | Crossing point of the CC potential | All parameters in this section are from Ursell 2007 (2) | 3 | nm |
| $a_{cross,OC}$ | Crossing point of the OC potential | | 2.75 | nm |
| $a_{cross,OO}$ | Crossing point of the OO potential | | 2.5 | nm |
| $E_{CC}$ | Value of the CC potential at $a_{min}$ | | -2.5 | $k_B T$ |
| $E_{OC}$ | Energy scale of the OC potential | | 50 | $k_B T$ |
| $E_{OO}$ | Value of the OO potential at $a_{min}$ | | -25 | $k_B T$ |
| $l_V$ | Potentials' decay length | | 1.5 | nm |

### OPEN PROBABILITY AND ITS DERIVATIVE WITH A LINEAR GROWTH OF THE CHANNEL POPULATION ACROSS DEVELOPMENT

**Figure 1:**
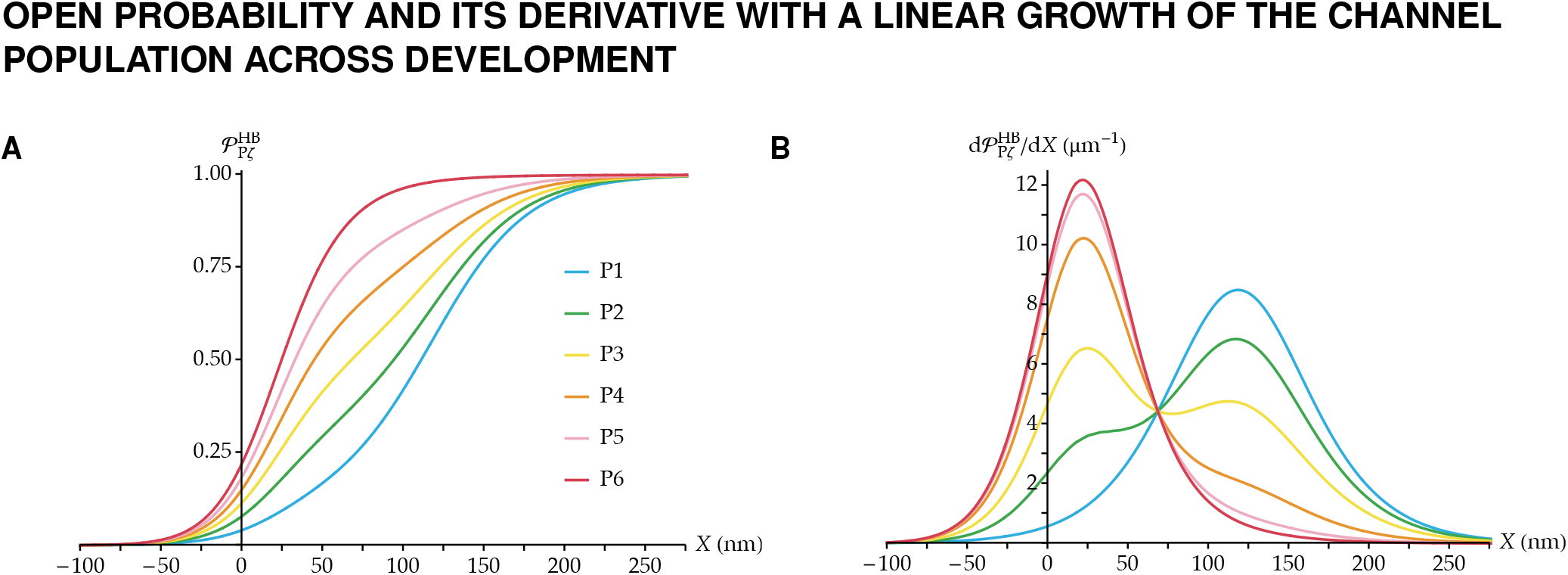
(A) Simulated maturation of the open-probability vs. displacement curve 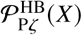 in a developing hair cell. We report the case of a linear growth of the number of MET channels over six days, as given in Table S1. The parameters that define the transduction units are the same as in Fig. 3 of the main text. (B) Derivatives of the open probability curves of panel *A*, using the same color code.

### MATURATION OF SLOW ADAPTATION

**Figure 2:**
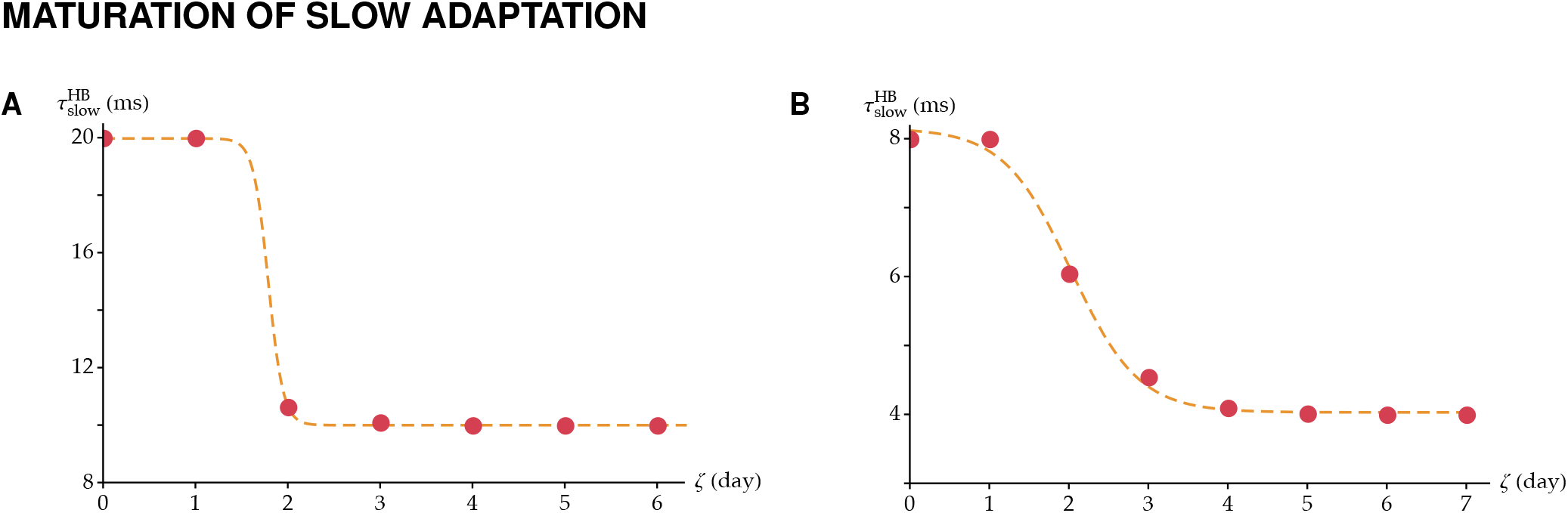
Time constant of slow adaptation 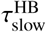 for the entire simulated hair bundle associated with the results reported in Fig. 6. (*A*) Slow adaptation time constant associated with Fig. 6A. (*B*) Slow adaptation time constant associated with Fig. 6C.

### EFFECT OF A SHIFT IN THE POPULATION GROWTH ON FAST ADAPTATION

**Figure 3:**
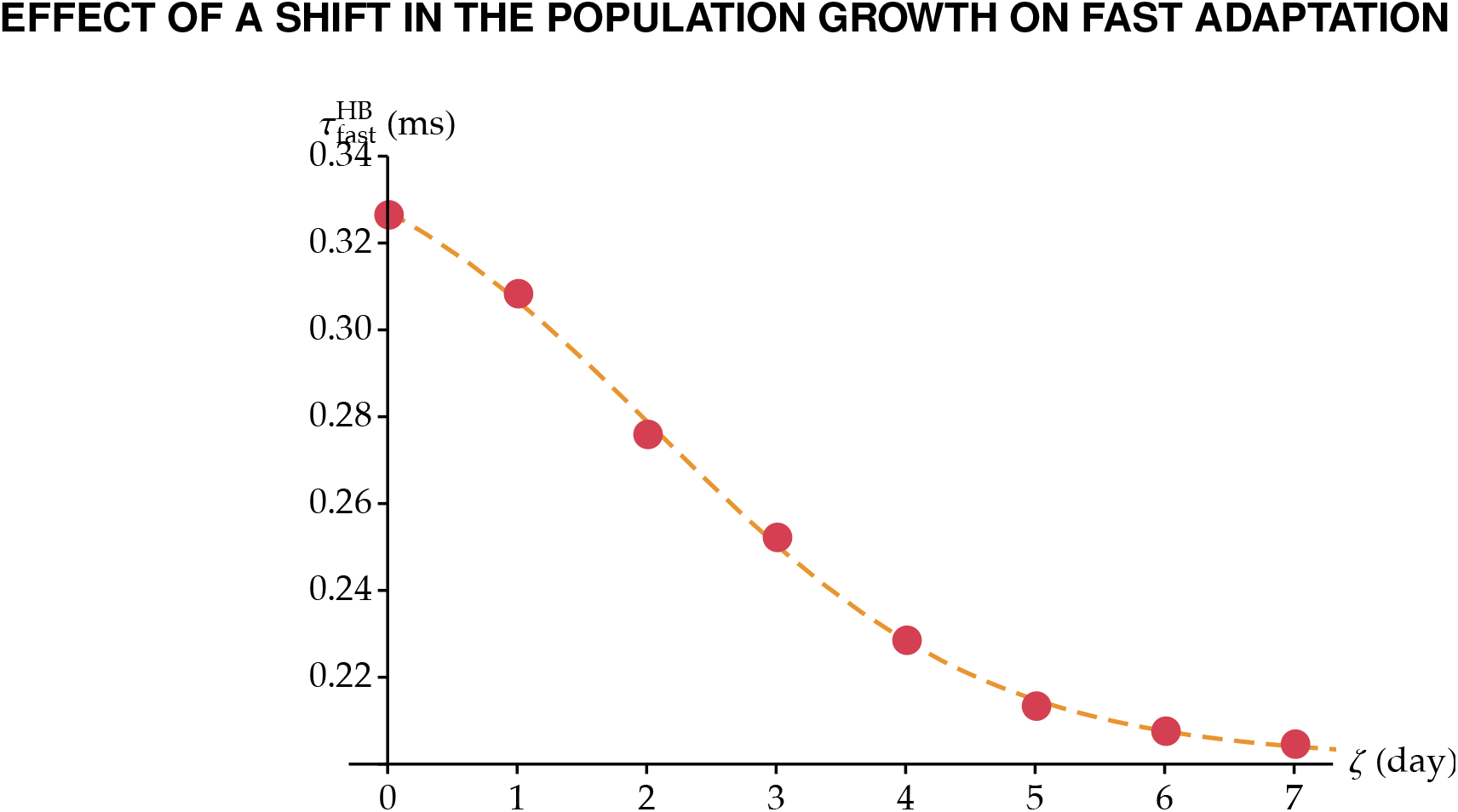
Simulated maturation of the time constant of fast adaptation 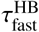 for the entire hair bundle as a function of developmental stages, assuming a sigmoidal growth in the number of channels over ten days, where the population growth has been shifted by three days to reach maturation at P7. Parameters are: 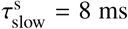, 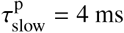, 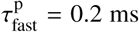, *a*^s^ = 0.7, and *a*^p^ = 0 as in Fig. 6*C,D*. A sigmoidal fit is used to visualize the trend.

### ADAPTATION KINETICS AS A FUNCTION OF THE TRANSDUCTION-CURRENT AMPLITUDE

**Figure 4:**
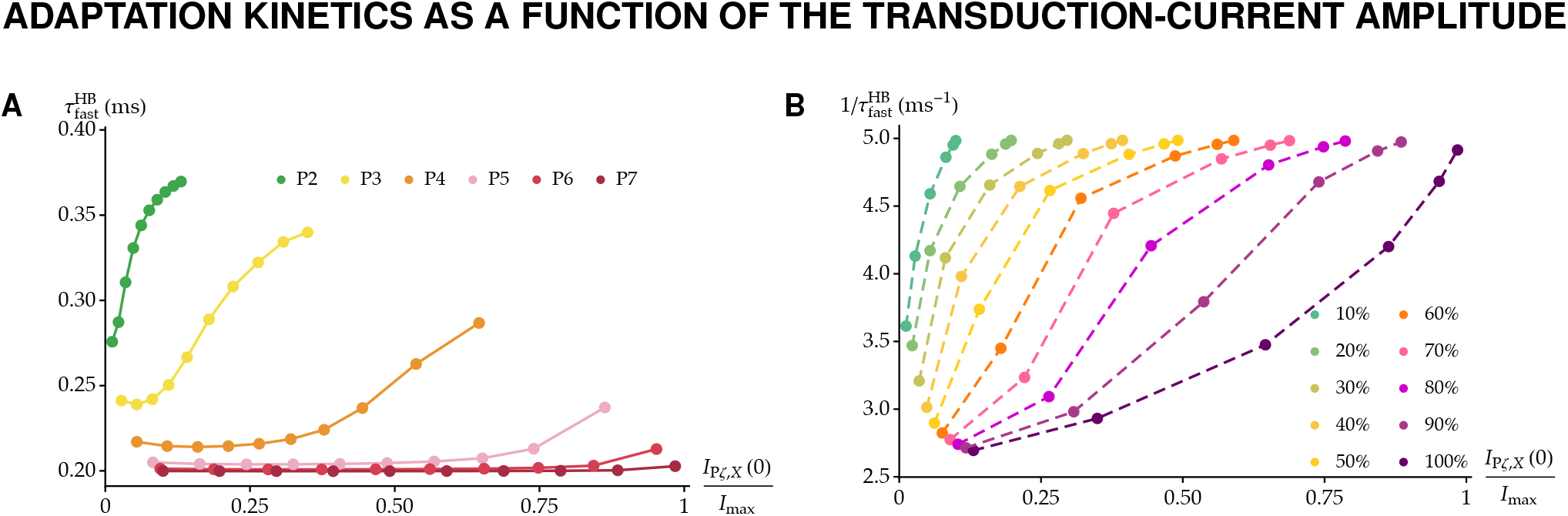
Adaptation kinetics as a function of the normalized peak transduction current at each developmental stage and imposed displacement, using the parameters determined from the data in Waguespack *et al.* (10) and Eq. 12 of the main text. (A) The time constant of fast adaptation for the whole hair bundle 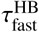 is plotted as a function of the normalized current at *t* = 0 as the displacement *X* is varied, at each developmental stage Pζ from P2 to P7. In each series, each data point corresponds to a different value of the overall open probability before adaptation 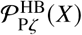, as given by Eq. 5 of the main text, from 10% to 100% in steps of 10%. (B) The rate of fast adaptation 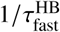 is plotted as a function of the same quantity as in (A), this time gathered per equal values of 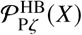. The plots are generated with the parameters characterizing the data in Waguespack *et al.* from the rat cochlea (10), as in Fig. 6C,D of the main text. Specifically, we assume a sigmoidal growth in the number of channels over seven days as reported in Table S1, and the other parameter values are: 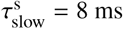, 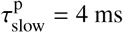, 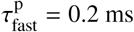, *a*^s^ = 0.7, and *a*^p^ = 0.

### COMPUTATION OF *X*_1/2_ IN THE GATING-SPRING MODEL

Within the standard gating-spring model (6), force balance at the level of the hair bundle reads (see Eq. 10 in (6)):

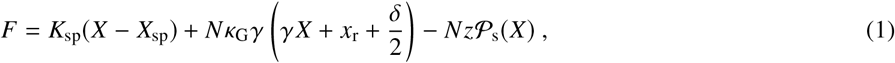

where *F* is the steady-state force required to hold the hair bundle at position *X*, *X*_sp_ is the resting position of the stereociliary pivots, *κ*_G_ is the gating-spring stiffness, *x*_r_ is the gating spring’s resting extension when the hair bundle is unperturbed, and all other parameters have been introduced in the main text and are summarized in Table S2. With our notation, Eqs. 6 and 8 of ref. (6) lead to *z* = *γk*_t_*δ*, as already reported in the main text, and *x*_r_ = (*E*_g_ − *zX*_1/2_)/(*κ*_G_*δ*). Imposing that the hair bundle sits at *X* = 0 when *F* = 0, we get, with *κ*_G_ = *k*_t_:

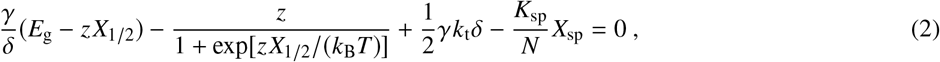

which specifies the value of *X*_1/2_.

## REFERENCES

1. Hudspeth, A. J., 1982. Extracellular current flow and the site of transduction by vertebrate hair cells. The Journal of Neuroscience 2:1–10.

2. Hudspeth, A. J., 1989. How the ear’s works work. Nature 341:397–404.

3. Assad, J. A., G. M. Shepherd, and D. P. Corey, 1991. Tip-link integrity and mechanical transduction in vertebrate hair cells. Neuron 7:985–994.

4. Howard, J., and A. J. Hudspeth, 1988. Compliance of the hair bundle associated with gating of mechanoelectrical transduction channels in the Bullfrog’s saccular hair cell. Neuron 1:189–199. www.sciencedirect.com/science/article/pii/0896627388901390.

5. Eatock, R. A., 2000. Adaptation in hair cells. Annual review of neuroscience 23:285–314. http://www.ncbi.nlm.nih.gov/pubmed/10845066.

6. Benser, M. E., R. E. Marquis, and A. J. Hudspeth, 1996. Rapid, Active Hair Bundle Movements in Hair Cells from the Bullfrog’s Sacculus. The Journal of Neuroscience 16:5629–5643. http://www.jneurosci.org/content/16/18/5629.

7. Ricci, A. J., A. C. Crawford, and R. Fettiplace, 2000. Active Hair Bundle Motion Linked to Fast Transducer Adaptation in Auditory Hair Cells. The Journal of Neuroscience 20:7131–7142. http://www.jneurosci.org/content/20/19/7131.long.

8. Corns, L. F., S. L. Johnson, C. J. Kros, and W. Marcotti, 2014. Calcium entry into stereocilia drives adaptation of the mechanoelectrical transducer current of mammalian cochlear hair cells. Proceedings of the National Academy of Sciences of the United States of America 111:14918–14923. http://www.pnas.org/cgi/doi/10.1073/pnas.1409920111.

9. Kennedy, H. J., M. G. Evans, A. C. Crawford, and R. Fettiplace, 2003. Fast adaptation of mechanoelectrical transducer channels in mammalian cochlear hair cells. Nature Neuroscience 6:832–836. http://www.nature.com.accesdistant.upmc.fr/neuro/journal/v6/n8/full/nn1089.html.

10. Ó Maoiléidigh, D., and A. J. Ricci, 2019. A Bundle of Mechanisms: Inner-Ear Hair-Cell Mechanotransduction. Trends in Neurosciences xx:1–16. https://linkinghub.elsevier.com/retrieve/pii/S0166223618303199.

11. Howard, J., and A. J. Hudspeth, 1987. Mechanical relaxation of the hair bundle mediates adaptation in mechanoelectrical transduction by the bullfrog’s saccular hair cell. Proceedings of the National Academy of Sciences of the United States of America 84:3064–3068. http://www.pubmedcentral.nih.gov/articlerender.fcgi?artid=304803{&}tool=pmcentrez{&}rendertype=abstract.

12. Si, F., H. Brodie, P. G. Gillespie, A. E. Vazquez, and E. N. Yamoah, 2003. Developmental assembly of transduction apparatus in chick basilar papilla. The Journal of Neuroscience 23:10815–10826.

13. Lelli, A., Y. Asai, A. Forge, J. R. Holt, and G. S. G. Geleoc, 2009. Tonotopic Gradient in the Developmental Acquisition of Sensory Transduction in Outer Hair Cells of the Mouse Cochlea. Journal of Neurophysiology 101:2961–2973. http://jn.physiology.org/cgi/oi/10.1152/jn.00136.2009.

14. Waguespack, J., F. T. Salles, B. Kachar, and A. J. Ricci, 2007. Stepwise Morphological and Functional Maturation of Mechanotransduction in Rat Outer Hair Cells. The Journal of Neuroscience 27:13890–13902. http://www.jneurosci.org/cgi/doi/10.1523/JNEUROSCI.2159-07.2007.

15. Indzhykulian, A. A., R. Stepanyan, A. Nelina, K. J. Spinelli, Z. M. Ahmed, I. A. Belyantseva, T. B. Friedman, P. G. Barr-Gillespie, and G. I. Frolenkov, 2013. Molecular Remodeling of Tip Links Underlies Mechanosensory Regeneration in Auditory Hair Cells. PLoS Biology 11:e1001583. http://dx.plos.org/10.1371/journal.pbio.1001583.

16. Ricci, A. J., Y. C. Wu, and R. Fettiplace, 1998. The endogenous calcium buffer and the time course of transducer adaptation in auditory hair cells. The Journal of Neuroscience 18:8261–77. http://www.ncbi.nlm.nih.gov/pubmed/9763471.

17. Wu, Y. C., A. J. Ricci, and R. Fettiplace, 1999. Two components of transducer adaptation in auditory hair cells. Journal of neurophysiology 82:2171–2181. http://jn.physiology.org/content/82/5/2171.

18. Zhao, Y.-D., E. N. Yamoah, and P. G. Gillespie, 1996. Regeneration of broken tip links and restoration of mechanical transduction in hair cells. Neurobiology Communicated by 94:15469–15474.

19. Lelli, A., P. Kazmierczak, Y. Kawashima, U. Müller, and J. R. Holt, 2010. Development and regeneration of sensory transduction in auditory hair cells requires functional interaction between cadherin-23 and protocadherin-15. The Journal of Neuroscience 30:11259–11269. http://www.jneurosci.org/cgi/reprint/30/34/11259.

20. Crawford, A. C., M. G. Evans, and R. Fettiplace, 1991. The actions of calcium on the mechano-electrical transducer current of turtle hair cells. The Journal of Physiology 434:369–398. http://jp.physoc.org/content/434/1/369.long{%}5Cnpapers2://publication/uuid/CA9D2A76-4B7E-4FEF-A564-BD51F605A6AE.

21. Cheung, E. L. M., and D. P. Corey, 2006. Ca2+ changes the force sensitivity of the hair-cell transduction channel. Biophysical journal 90:124–39. http://www.pubmedcentral.nih.gov/articlerender.fcgi?artid=1367012{&}tool=pmcentrez{&}rendertype=abstract{%}5Cn.

22. Kim, K. X., M. Beurg, C. M. Hackney, D. N. Furness, S. Mahendrasingam, and R. Fettiplace, 2013. The role of transmembrane channel–like proteins in the operation of hair cell mechanotransducer channels. The Journal of General Physiology 142:493–505. http://www.jgp.org/lookup/doi/10.1085/jgp.201311068.

23. Corns, L. F., J.-Y. Jeng, G. P. Richardson, C. J. Kros, and W. Marcotti, 2017. Tmc2 modifies permeation properties of the mechanoelectrical transducer channel in early postnatal mouse cochlear outer hair cells. Frontiers in Molecular Neuroscience 10:1–15. http://journal.frontiersin.org/article/10.3389/fnmol.2017.00326/full.

24. Kurima, K., L. M. Peters, Y. Yang, S. Riazuddin, Z. M. Ahmed, S. Naz, D. Arnaud, S. Drury, J. Mo, T. Makishima, M. Ghosh, P. S. Menon, D. Deshmukh, C. Oddoux, H. Ostrer, S. Khan, S. Riazuddin, P. L. Deininger, L. L. Hampton, S. L. Sullivan, J. F. Battey, B. J. Keats, E. R. Wilcox, T. B. Friedman, and A. J. Griffith, 2002. Dominant and recessive deafness caused by mutations of a novel gene, TMC1, required for cochlear hair-cell function. Nature Genetics 30:277–284.

25. Kawashima, Y., G. S. G. Géléoc, K. Kurima, V. Labay, A. Lelli, Y. Asai, T. Makishima, D. K. Wu, C. C. D. Santina, J. R. Holt, and A. J. Griffith, 2011. Mechanotransduction in mouse inner ear hair cells requires transmembrane channel-like genes. Journal of Clinical Investigation 121:4796–4809.

26. Kurima, K., S. Ebrahim, B. Pan, M. Sedlacek, P. Sengupta, B. A. Millis, R. Cui, H. Nakanishi, T. Fujikawa, Y. Kawashima, B. Y. Choi, K. Monahan, J. R. Holt, A. J. Griffith, and B. Kachar, 2015. TMC1 and TMC2 Localize at the Site of Mechanotransduction in Mammalian Inner Ear Hair Cell Stereocilia. Cell Reports 12:1606–1617. http://dx.doi.org/10.1016/j.celrep.2015.07.058.

27. Mahendrasingam, S., and D. N. Furness, 2019. Ultrastructural localization of the likely mechanoelectrical transduction channel protein, transmembrane-like channel 1 (TMC1) during development of cochlear hair cells. Scientific Reports 9:1–9. http://dx.doi.org/10.1038/s41598-018-37563-x.

28. Beurg, M., R. Fettiplace, J.-h. Nam, and A. J. Ricci, 2009. Localization of inner hair cell mechanotransducer channels using high speed calcium imaging. Nature neuroscience 12:553–558.

29. Beurg, M., R. Cui, A. C. Goldring, S. Ebrahim, R. Fettiplace, and B. Kachar, 2018. Variable number of TMC1-dependent mechanotransducer channels underlie tonotopic conductance gradients in the cochlea. Nature Communications http://dx.doi.org/10.1038/s41467-018-04589-8.

30. Gianoli, F., T. Risler, and A. S. Kozlov, 2017. Lipid bilayer mediates ion-channel cooperativity in a model of hair-cell mechanotransduction. Proceedings of the National Academy of Sciences of the United States of America 114:E11010–E11019. http://www.pnas.org/lookup/doi/10.1073/pnas.1713135114.

31. Feller, W., 1968. An Introduction to Probability Theory and Its Applications −Vol. I. Wiley.

32. Cotton, J., and W. Grant, 2004. Computational models of hair cell bundle mechanics: II. Simplified bundle models. Hearing Research 197:105–111.

33. Kozlov, A. S., T. Risler, and A. J. Hudspeth, 2007. Coherent motion of stereocilia assures the concerted gating of hair-cell transduction channels. Nature neuroscience 10:87–92. http://www.pubmedcentral.nih.gov/articlerender.fcgi?artid=2174432{&}tool=pmcentrez{&}rendertype=abstract.

34. Karavitaki, K. D., and D. P. Corey, 2010. Sliding Adhesion Confers Coherent Motion to Hair Cell Stereocilia and Parallel Gating to Transduction Channels. The Journal of Neuroscience 30:9051–9063. http://www.jneurosci.org/cgi/doi/10.1523/JNEUROSCI.4864-09.2010.

35. Kozlov, A. S., J. Baumgart, T. Risler, C. P. C. Versteegh, and A. J. Hudspeth, 2011. Forces between clustered stereocilia minimize friction in the ear on a subnanometre scale. Nature 474:376–379. http://www.nature.com/doifinder/10.1038/nature10073.

36. Kozlov, A. S., T. Risler, A. J. Hinterwirth, and A. Hudspeth, 2012. Relative stereociliary motion in a hair bundle opposes amplification at distortion frequencies. The Journal of Physiology 590:301–308. http://doi.wiley.com/10.1113/jphysiol.2011.218362.

37. Beurg, M., M. G. Evans, C. M. Hackney, and R. Fettiplace, 2006. A large-conductance calcium-selective mechanotransducer channel in mammalian cochlear hair cells. The Journal of Neuroscience 26:10992–11000. http://www.jneurosci.org/cgi/doi/10.1523/JNEUROSCI.2188-06.2006.

38. Peng, A. W., F. T. Salles, B. Pan, and A. J. Ricci, 2011. Integrating the biophysical and molecular mechanisms of auditory hair cell mechanotransduction. Nature Communications 2:523. http://dx.doi.org/10.1038/ncomms1533.

39. Fettiplace, R., and K. X. Kim, 2014. The Physiology of Mechanoelectrical Transduction Channels in Hearing. Physiological Reviews 94:951–986. http://physrev.physiology.org/cgi/doi/10.1152/physrev.00038.2013.

40. Peng, A. W., and A. J. Ricci, 2009. Auditory and Vestibular Research. Methods in molecular biology (Clifton, N.J.) 493:487–500. http://link.springer.com/10.1007/978-1-59745-523-7.

41. Tobin, M., A. Chaiyasitdhi, V. Michel, N. Michalski, and P. Martin, 2019. Stiffness and tension gradients of the hair cell’s tip-link complex in the mammalian cochlea. eLife 8:1–36.

42. Ursell, T., K. C. Huang, E. Peterson, and R. Phillips, 2007. Cooperative gating and spatial organization of membrane proteins through elastic interactions. PLoS Computational Biology 3:0803–0812. http://journals.plos.org/ploscompbiol/article?id=10.1371/journal.pcbi.0030081.

43. Bean, B. P., 1989. Neurotransmitter inhibition of neuronal calcium currents by changes in channel voltage dependence. Nature 340:153–156.

44. Marcotti, W., L. F. Corns, R. J. Goodyear, A. K. Rzadzinska, K. B. Avraham, K. P. Steel, G. P. Richardson, and C. J. Kros, 2016. The acquisition of mechano-electrical transducer current adaptation in auditory hair cells requires myosin VI. Journal of Physiology 594:3667–3681.

45. Husbands, J. M., S. A. Steinberg, R. Kurian, and J. C. Saunders, 1999. Tip-link integrity on chick tall hair cell stereocilia following intense sound exposure. Hearing Research 135:135–145.

46. Vollrath, M. A., and R. A. Eatock, 2003. Time course and extent of mechanotransducer adaptation in mouse utricular hair cells: comparison with frog saccular hair cells. Journal of neurophysiology 90:2676–89. http://www.ncbi.nlm.nih.gov/pubmed/12826658.

47. Stauffer, E. A., J. D. Scarborough, M. Hirono, E. D. Miller, K. Shah, J. A. Mercer, J. R. Holt, and P. G. Gillespie, 2005. Fast adaptation in vestibular hair cells requires Myosin-1c activity. Neuron 47:541–553.

48. Pan, B., G. S. Géléoc, Y. Asai, G. C. Horwitz, K. Kurima, K. Ishikawa, Y. Kawashima, A. J. Griffith, and J. R. Holt, 2013. TMC1 and TMC2 are components of the mechanotransduction channel in hair cells of the mammalian inner ear. Neuron 79:504–515. http://www.sciencedirect.com/science/article/pii/S0896627313005357.

49. Beurg, M., J.-H. Nam, Q. Chen, and R. Fettiplace, 2010. Calcium Balance and Mechanotransduction in Rat Cochlear Hair Cells. Journal of Neurophysiology 104:18–34. http://jn.physiology.org/cgi/doi/10.1152/jn.00019.2010.

50. Choe, Y., M. O. Magnasco, and A. J. Hudspeth, 1998. A model for amplification of hair-bundle motion by cyclical binding of Ca2+ to mechanoelectrical-transduction channels. Proceedings of the National Academy of Sciences of the United States of America 95:15321–15326. http://www.pnas.org/content/95/26/15321.

51. Kozlov, A. S., D. Andor-Ardo, and A. J. Hudspeth, 2012. Anomalous Brownian motion discloses viscoelasticity in the ear’s mechanoelectrical-transduction apparatus. Proceedings of the National Academy of Sciences 109:2896–2901.

52. Bartsch, T. F., F. E. Hengel, A. Oswald, G. Dionne, I. V. Chipendo, S. S. Mangat, M. E. Shatanofy, L. Shapiro, U. Müller, and A. J. Hudspeth, 2019. Elasticity of Individual Protocadherin 15 Molecules Implicates Tip Links as the Gating Springs for Hearing. Proceedings of the National Academy of Sciences 201902163.

53. Corns, L. F., S. L. Johnson, T. Roberts, K. M. Ranatunga, A. Hendry, F. Ceriani, S. Safieddine, K. P. Steel, A. Forge, C. Petit, D. N. Furness, C. J. Kros, and W. Marcotti, 2018. Mechanotransduction is required for establishing and maintaining mature inner hair cells and regulating efferent innervation. Nature Communications 9:4015. https://doi.org/10.1038/s41467-018-06307-w.

## REFERENCES

1. Tobin, M., A. Chaiyasitdhi, V. Michel, N. Michalski, and P. Martin, 2019. Stiffness and tension gradients of the hair cell’s tip-link complex in the mammalian cochlea. eLife 8:1–36.

2. Ursell, T., K. C. Huang, E. Peterson, and R. Phillips, 2007. Cooperative gating and spatial organization of membrane proteins through elastic interactions. PLoS Computational Biology 3:0803–0812. http://journals.plos.org/ploscompbiol/article?id=10.1371/journal.pcbi.0030081.

3. Corey, D. P., and A. J. Hudspeth, 1983. Analysis of the microphonic potential of the bullfrog’s sacculus. The Journal of neuroscience 3:942–961. http://eutils.ncbi.nlm.nih.gov/entrez/eutils/elink.fcgi?dbfrom=pubmed{&}id=6601693{&}retmode=ref{&}cmd=prlinks{%}5Cnpapers2://publication/uuid/22C5E385-C31B-4371-89B3-ADD282A4770F.

4. Hudspeth, A. J., 1992. Hair-bundle mechanics and a model for mechanoelectrical transduction by hair cells. Soc Gen Physiol Ser 47:357–370. http://www.ncbi.nlm.nih.gov/entrez/query.fcgi?cmd=Retrieve{&}db=PubMed{&}dopt=Citation{&}list{_}uids=1369770.

5. Ricci, A. J., B. Kachar, J. Gale, and S. M. Van Netten, 2006. Mechano-electrical transduction: New insights into old ideas. Journal of Membrane Biology 209:71–88. http://link.springer.com.accesdistant.upmc.fr/article/10.1007/s00232-005-0834-8.

6. Howard, J., and A. J. Hudspeth, 1988. Compliance of the hair bundle associated with gating of mechanoelectrical transduction channels in the Bullfrog’s saccular hair cell. Neuron 1:189–199. http://www.sciencedirect.com/science/article/pii/0896627388901390.

7. Kachar, B., M. Parakkal, M. Kurc, Y.-d. Zhao, and P. G. Gillespie, 2000. High-resolution structure of hair-cell tip links. Proceedings of the National Academy of Sciences of the United States of America 97:13336–41. http://www.pubmedcentral.nih.gov/articlerender.fcgi?artid=27225{&}tool=pmcentrez{&}rendertype=abstract.

8. Araya-Secchi, R., B. L. Neel, and R. Sotomayor, 2016. An Elastic Element in the Protocadherin-15 Tip Link of the Inner Ear. Nature communications 7:1–14. http://dx.doi.org/10.1038/ncomms13458.

9. Powers, R. J., S. Roy, E. Atilgan, W. E. Brownell, S. X. Sun, P. G. Gillespie, and A. A. Spector, 2012. Stereocilia membrane deformation: Implications for the gating spring and mechanotransduction channel. Biophysical Journal 102:201–210. http://www.sciencedirect.com/science/article/pii/S0006349511054245.

10. Waguespack, J., F. T. Salles, B. Kachar, and A. J. Ricci, 2007. Stepwise Morphological and Functional Maturation of Mechanotransduction in Rat Outer Hair Cells. The Journal of Neuroscience 27:13890–13902. http://www.jneurosci.org/cgi/doi/10.1523/JNEUROSCI.2159-07.2007.

